# TIP5 safeguards genome architecture of ground-state pluripotent stem cells

**DOI:** 10.1101/2019.12.20.882282

**Authors:** Damian Dalcher, Jennifer Yihong Tan, Cristiana Bersaglieri, Rodrigo Peña-Hernández, Eva Vollenweider, Stefan Zeyen, Marc W. Schmid, Valerio Bianchi, Stefan Butz, Rostyslav Kuzyakiv, Tuncay Baubec, Ana Claudia Marques, Raffaella Santoro

## Abstract

Chromosomes have an intrinsic tendency to segregate into compartments, forming long-distance contacts between loci of similar chromatin states. However, how genome compartmentalization is regulated remains elusive. We analyzed two closely and developmentally related pluripotent cell types: ground-state ESCs that have an open and active chromatin and developmentally advanced ESCs that display a more closed and repressed state. We show that these two ESC types differ in their regulation of genome organization due to their differential dependency on TIP5, a component of the chromatin remodeling complex NoRC. We show that TIP5 interacts on ESC chromatin with SNF2H, DNA topoisomerase 2A (TOP2A) and cohesin. TIP5 associates with sub-domains within the active A compartment that strongly intersect through long-range contacts in ESCs. We found that only ground-state chromatin requires TIP5 to limit the invasion of active domains into repressive compartments. Depletion of TIP5 increased chromatin accessibility particularly at B compartments and decreased their repressive features. Furthermore, TIP5 acts as a barrier for the repressive H3K27me3 spreading, a process that also requires TOP2A activity. Finally, ground-state ESCs require TIP5 for growth, differentiation capacity, and correct expression of developmental genes. Our results revealed the propensity of open and active chromatin domains to invade repressive domains, an action counteracted by chromatin remodeling and the relief of chromatin torsional stress. This effort in controlling open/active chromatin domains is required to establish active and repressed genome partitioning and preserves cell function and identity.

## Introduction

The 3D genome organization is functionally important for correct execution of gene expression programs. Chromosomes have an intrinsic tendency to segregate into compartments based on the local epigenetic landscape and transcriptional activity. At the megabase scale, the genome can be divided into two compartments, called A and B compartments, which are estimated by an eigenvector analysis of the genome contact matrix after normalization by the observed-expected method ^1^. Genomic contacts predominately occur between loci belonging to the same compartment. Compartment A is highly enriched for open and active chromatin whereas compartment B is enriched for closed and repressed chromatin. Furthermore, long-distance contacts have been shown to occur between domains with similar chromatin states ^2^. Removal or depletion of chromatin-associated cohesin eliminates all loop domains and increased compartmentalization ^3–6^. The opposite effect was achieved by increasing the residence time of cohesins on DNA, which leads to extension of chromatin loops and a less strict segregation between A and B compartments with a decrease of far-*cis* contacts^5^. In contrast to topologically associating domains (TAD) boundaries, which are generally conserved among cell types, A and B compartments are cell type specific ^2, 7, 8^. However, the mechanisms underlying these changes and their impact on gene expression and epigenetic states remain yet unclear.

Pluripotent embryonic stem cells (ESCs) provide an exceptional system to address this question as they exist in a variety of states that represent distinct chromatin and epigenetic features. *In vitro*, ESC types are largely defined by culture conditions, which mimic the natural development of the embryo from the blastocyst to post-implantation stages ^9^. ESCs can be propagated in medium containing fetal calf serum and leukaemia inhibitory factor (LIF) (ESC+serum) or in serum-free 2i medium (ESC+2i) that contains LIF plus two small-molecule kinase inhibitors for MEK/ERK (PD0325901) and GSK3 (CHIR99021) ^10^. Both ESC+2i and ESC+serum are pluripotent. ESC+2i closely resemble the developmental ground-state *in vivo* whereas ESC+serum are developmentally advanced ^11^. These two pluripotent cell types display distinct chromatin and epigenetic features. At the nucleosomal level, the chromatin configuration is more open in ESC+2i than in ESC+serum ^12^. Furthermore, the epigenetic and chromatin organization of ESC+2i is in a less repressed state. ESC+2i have hypomethylated DNA similar to pre-implantation embryos, whereas ESC+serum genome is hypermethylated, reminiscent of post-implantation embryos ^13–16^. Similarly, there is a reduced prevalence of the repressive H3K27me3 mark at Polycomb targets in ESC+2i ^16, 17^. Recent studies showed that ESC+serum contain a set of extremely long-range interactions that depend on Polycomb repressive complexes 1 and 2 (PRC1, PRC2) ^17, 18^. In contrast, ESC+2i do not contain these long-range PRC interactions and instead exhibit chromatin decompaction at PRC target loci ^17, 19^.

In this work, we made use of these two closely and developmentally related ESC types to elucidate how genome organization and compartmentalization are regulated according to cell state and chromatin structure. We show that only the active and open genome of ground-state ESCs rely on TIP5, a factor that together with SNF2H constitutes the nucleolar remodelling complex NoRC^20^. TIP5 interacts on ESC chromatin with SNF2H, DNA topoisomerase 2A (TOP2A) and cohesin and associates with active sub-domains within A compartments. We found that the strongest long-distance contacts in ESCs are between TIP5-bound regions. TIP5 specifically regulates the highly open chromatin of ESC+2i by limiting the invasion of active domains into repressive compartments. Depletion of TIP5 specifically affects ground-state ESCs by increasing chromatin accessibility at chromatin regions including the repressed B compartments, which in turn decrease their repressive features such as upregulation of gene expression and acquisition of active epigenetic marks. Furthermore, the binding of TIP5 with active chromatin domains of ground-state ESCs acts as a barrier for H3K27me3 spreading, a process that also involves TOP2A activity. Finally, the ground-state specific role of TIP5 was also evident by defects in proliferation, impaired differentiation capacity, and deregulation of genes linked to developmental process, which all occur only upon TIP5 depletion in ESC+2i whereas ESC+serum remained unaffected. Our results suggest that chromatin remodeling and the relief of chromatin torsional stress might serve to counteract the intrinsic propensity of the highly open and active chromatin of ground-state ESCs to invade repressive domains. This effort in controlling open/active chromatin domains is required to establish active and repressed genome partitioning and preserves cell function and identity.

## Results

### Ground-state and developmentally advanced pluripotent ESCs differ in the requirement of TIP5 for cell proliferation and differentiation capacity

The nucleolar remodelling complex NoRC consists of two subunits, the ATPase SMARCA5/SNF2H and TIP5, a >200 kDa protein that shares sequence homology with the largest subunits of SNF2H/ISWI-containing remodeling complexes ^20, 21^. In differentiated cells, TIP5 is mainly localized in nucleoli, associates with ribosomal RNA (rRNA) genes and establishes their epigenetic silencing ^20, 22, 23^. In contrast, in ESCs, recruitment of TIP5 to rRNA genes and its ability to silence rRNA genes is impaired through a long non-coding RNA-mediated mechanism ^24, 25^.

Our previous work and initial analysis suggested that in ESCs TIP5 function was not related to rRNA transcriptional control and was distinct between ground-state and developmentally advanced ESCs. First, TIP5 is more highly expressed in ESC+2i than in differentiated cells (**Fig. 1a**). Second, although not bound to rRNA genes, TIP5 is still tightly associated with chromatin in ESC+2i (**Fig. 1b**). Third, proliferation of ESC+2i decreased upon TIP5 depletion by siRNA (**Fig. 1c,d**). Similar results were obtained with a different siRNA and in another ESC line (**Supplementary Figure 1a,b**). TIP5 downregulation in ESC+2i induced a moderate arrest at G1 phase of cell cycle without any evident sign of apoptotic cell death (**Supplementary Figure 1c-e**). Fourth, and consistent with previous results ^24^, after induction of monolayer differentiation upon withdrawal of LIF, ESC+2i treated with siRNA-*Tip5* underwent cell death while control cells displayed morphological structures typical of differentiated cells and were negative for alkaline phosphatase staining (**Fig. 1f,g**). All these results indicated that ground-state ESCs depend on TIP5 expression for proliferation and differentiation capacity and highlighted an unexpected non-nucleolar function of TIP5. Next we asked whether proliferation and differentiation capacities of developmentally advanced ESC+serum were also dependent on TIP5. Surprisingly, although TIP5 expression levels and knockdown efficiency in ESC+2i and ESC+serum were similar (**Fig. 1c,e**), depletion of TIP5 in ESC+serum did not cause any evident defect in proliferation or differentiation (**Fig. 1d,f,g**), suggesting ground-state specific role of TIP5. To further support these results, we performed CRIPR/Cas9 to generate TIP5-KO directly in ESC+serum and ESC+2i. We were able to generate TIP5-KO lines only in ESC+serum (**Fig. 1h**). whereas all our attempts to establish TIP5-KO directly in ESC+2i failed, a result that is consistent with the proliferative defects observed upon siRNA-mediated TIP5 depletion only in ESC+2i, but not in ESC+serum (**Fig. 1h**). Cell morphology, proliferation, and differentiation capacity are similar between control and TIP5-KO ESC+serum (**Fig. 1i,j, Supplementary Figure 1f**). However, during the transition from serum to 2i conditions TIP5-KO cells lose self-renewal and proliferation capacity with substantial cell death (**Fig. 1j, Supplementary Figure 1g**), indicating that TIP5 is specifically essential in ground-state ESCs.

**Figure 1.**
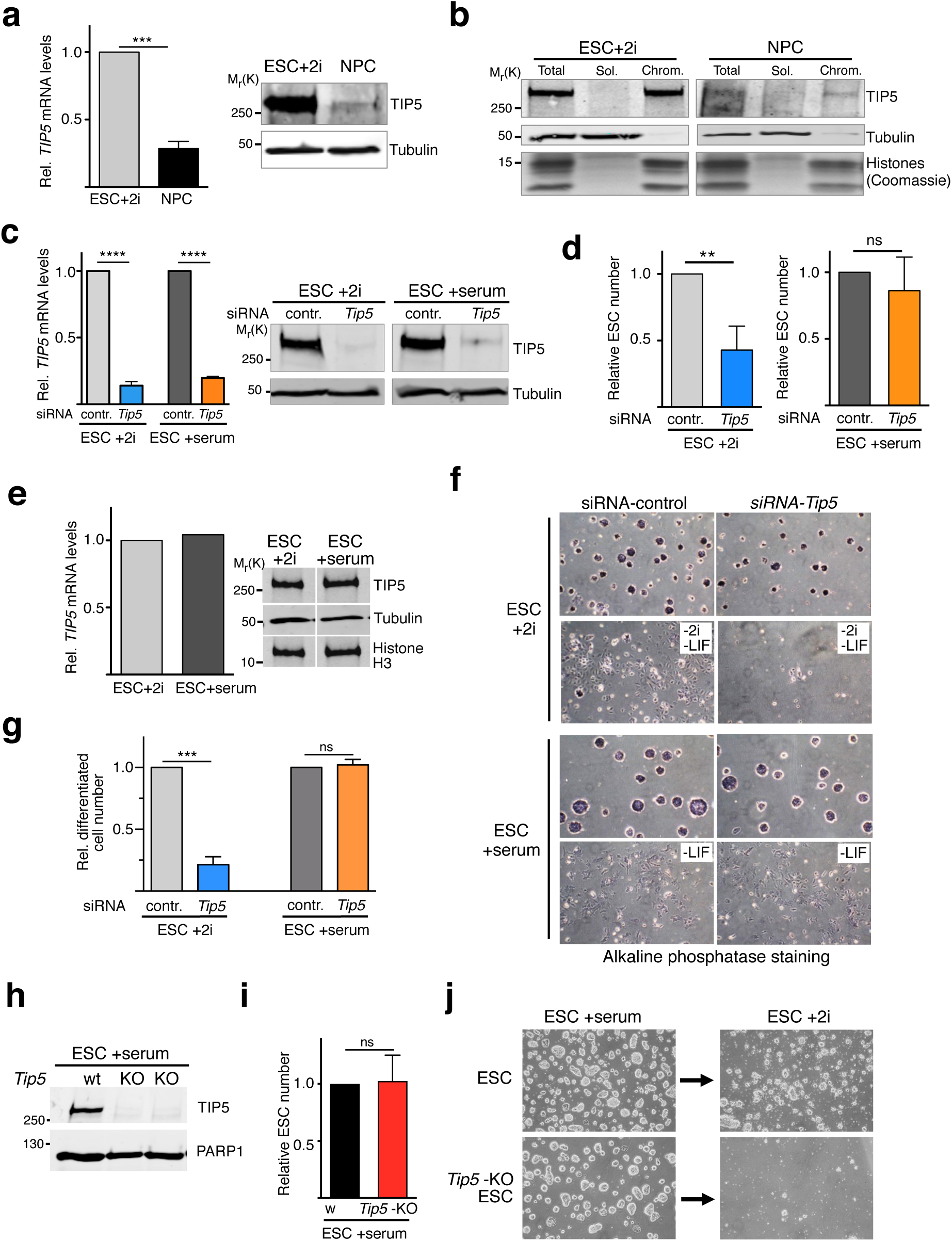
TIP5 is required for proliferation and differentiation of ESC+2i. **a.** TIP5 expression is higher in ESC+2i than in differentiated cells (neural progenitors, NPC). Left panel. *TIP5* mRNA levels were measured by qRT-PCR and normalized to *Rps12* mRNA and to ESC+2i. Average values of three independent experiments. Error bars represent s.d. and statistical significance (*P*-values) was calculated using the paired two-tailed t-test (***< 0.001).Right panel. Western blot showing TIP5 protein levels in ESC+2i and NPC. Tubulin is shown as a protein loading control. **b.** TIP5 associates with chromatin of ESC and NPC. Chromatin-bound (Chrom.) and soluble (Sol.) fractions of equivalent cell number of ESC+2i and NPCs were analyzed by western blot for TIP5 levels. Tot., total protein. Tubulin and histones are shown as loading and fractionation control. **c.** siRNA-knockdown efficiency of TIP5 shown by qRT-PCR and western blot. *TIP5* mRNA levels were measured by qRT-PCR and normalized to *Rps12* mRNA and to each ESC line. Average values of three independent experiments. Error bars represent s.d. Error bars represent s.d. and statistical significance (*P*-values) was calculated using the paired two-tailed t-test (****< 0.0001).Right panel. Western blot showing TIP5 protein levels. Tubulin is shown as a protein loading control. **d.** TIP5 knockdown affects proliferation of ESC+2i but not of ESC+serum. Data represent relative cell numbers after 3 days of siRNA treatment and were normalized to ESC transfected with siRNA-Control. Average values of three independent experiments. Error bars represent s.d. Statistical significance (*P*-values) for the experiments was calculated using the paired two-tailed t-test (** < 0.01; ns, non-significant). **e.** TIP5 is expressed at similar levels in both ESC+2i and ESC+serum. Left panel. *TIP5* mRNA levels were measured by qRT-PCR and normalized to *Rps12* mRNA and to ESC+2i. Average values of three independent experiments. Error bars represent s.d.. Right panel. Western blot showing TIP5 protein levels in ESC+2i and ESC+serum. Tubulin and histone H3 are shown as a protein loading controls. **f.** TIP5 is required for the differentiation of ESC+2i but not of ESC+serum. Representative images of alkaline phosphatase staining of ESC and cells after 3 days of differentiation. **g.** Quantification of differentiated cells. Values represent relative number of differentiated cells from three independent experiments relative to control cells. Error bars represent s.d. Statistical significance (*P*-values) for the experiments was calculated using the paired two-tailed t-test (*** < 0.001; ns, non-significant). **h.** Western blot showing TIP5 protein levels in two TIP5-KO ESC lines obtained via CRISPr/Cas9 directly in ESC+serum. PARP1 is shown as a protein loading control. **i.** Cell proliferation is not affected in TIP5-KO ESC+serum. Data represent relative cell numbers 2 days culture starting with the same number of cells. Average values of four independent experiments. Error bars represent s.d. Statistical significance (*P*-values) for the experiments was calculated using the paired two-tailed t-test (ns, non-significant). **i.** Representative images of wt and *Tip5*-KO ESC+serum before and after transition in 2i conditions.

Taken together these results highlighted a substantial difference in the requirement of TIP5 for cell proliferation and differentiation capacities between ground-state and developmentally advanced pluripotent ESCs.

### TIP5 regulates gene expression in ground-state ESCs

To elucidate the ground-state specific role of TIP5, we analysed gene expression of ESC+2i and ESC+serum treated with siRNA-*Tip5* or siRNA-control (**Fig. 2a, Supplementary Table 1**). We found that depletion of TIP5 induces greater differential gene expression in ESC+2i than in ESC+serum. Upon TIP5 depletion in ESC+2i, 1934 genes showed transcriptional changes (log_2_ fold change > 0.58; *P*<0.05; 1236 upregulated and 698 downregulated). In contrast, ESC+serum depleted for TIP5 showed only moderate changes in gene expression compared to ESC+2i, and the total number of genes affected by TIP5 knockdown was almost 4 times less (351 upregulated and 207 downregulated) (**Fig. 2a**). Only a minority of differentially expressed genes were upregulated (65) or downregulated (56) by TIP5 knockdown in both ESC states (**Supplementary Figure 2a**). Validation by qRT-PCR, using 2 siRNAs and a different ESC line, supported a role of TIP5 in the regulation of gene expression in ESC+2i but not in ESC+serum (**Fig. 2b, Supplementary Figure 2b,c**). Gene ontology analysis revealed that differentially expressed genes in ESC+2i are enriched in pathways linked to developmental processes (**Fig. 2c, Supplementary Table 2**). In contrast, genes differentially expressed in ESC+serum are mostly implicated in biological processes linked to cell signalling and cell adhesion.

**Figure 2.**
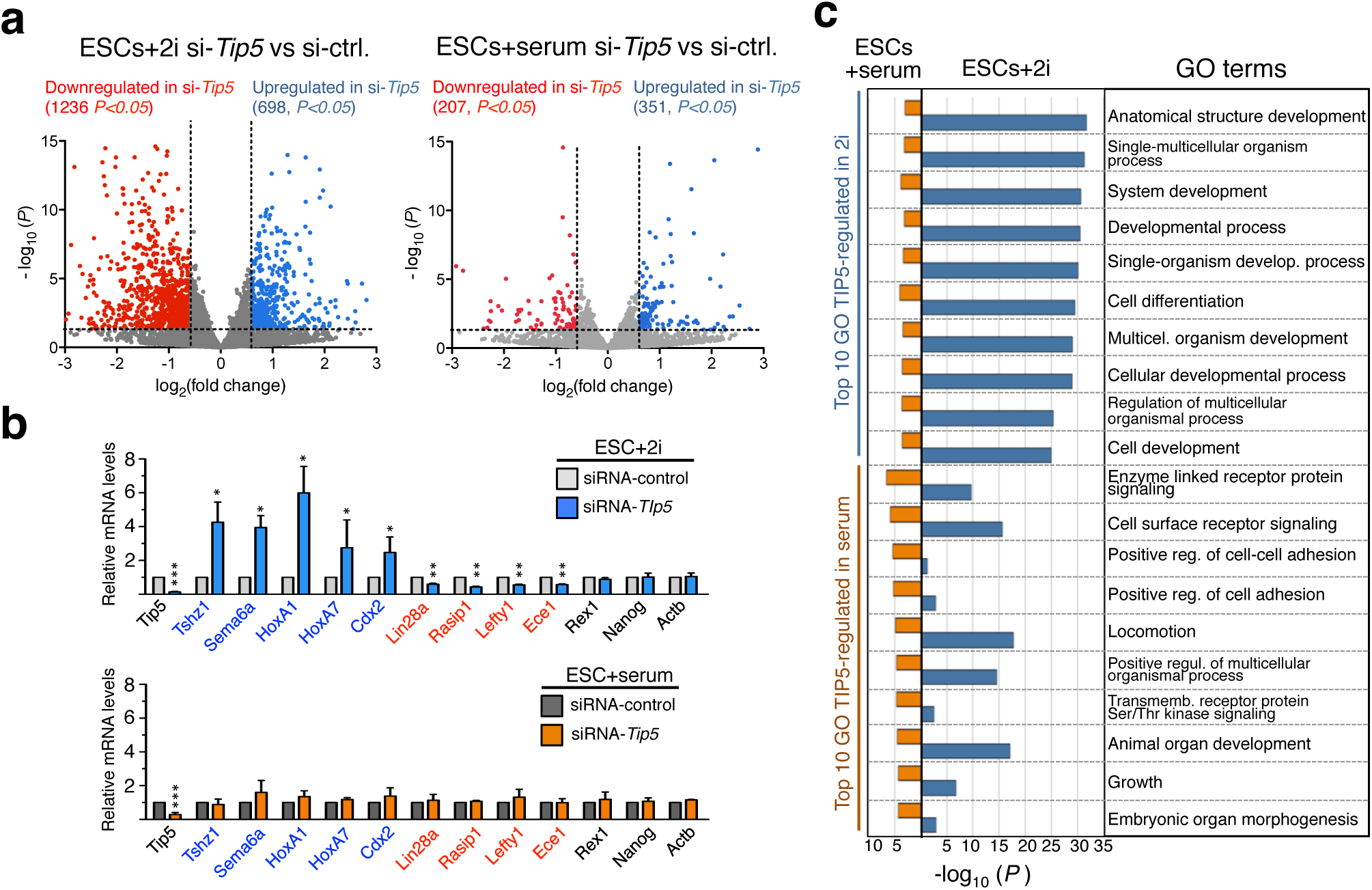
TIP5 regulate gene expression specifically in ESC+2i. **a.** Volcano plot showing fold change (log_2_ values) in transcript level of ESC+2i and ESC+serum upon TIP5 knockdown. Gene expression values of three replicates were averaged and selected for 1.5 fold changes and *P* <0.05. **b.** Validation by qRT-PCR of genes regulated by TIP5 in ESC+2i but not in ESC+serum (upregulated genes in ESC+2i upon TIP5-KD are labeled in blue, downregulated genes are in red). *Nanog*, *Rex1* and *Actin* B (*ActB*) are shown as genes not regulated by TIP5. mRNA levels were normalized to *Rps12* mRNA and to ESCs transfected with siRNA-Control. Average values of three independent experiments. Error bars represent s.d. Statistical significance (*P*-values) for the experiments was calculated using the paired two-tailed t-test (*<0.05, **< 0.01, ***<0.001). Data without P values are statistically non significant. **c.** Top 10 biological process gene ontology terms as determined using DAVID for genes regulated by TIP5 in ESC+2i and ESC+serum.

Next, we asked whether the role of TIP5 in the regulation of gene expression specifically in ESC+2i was a consequence of the chromatin state of these cells (low DNA methylation and low H3K27me3 occupancy at PRC targets). To mimic this status in ESC+serum, we analysed ESCs that lack DNA methyltransferases DNMT1, 3B and 3A (TKO-ESC) or the core components of PRC1 and PRC2 complexes (Ring1b^−/−^ ESC and Eed^−/−^ ESC). TIP5 depletion in these ESC-KO lines cultured in serum did not affect cell proliferation or expression of TIP5 2i-regulated genes, indicating that the specific function of TIP5 in ESC+2i does not directly depend on low DNA methylation and H327me3 content (**Supplementary Figures 2d-g**). Given the higher H3K27me3 content at Polycomb targets in ESC+serum^16^, genes regulated by TIP5 in 2i also display higher H3K27me3 occupancies in ESC+serum than in ESC+2i (**Supplementary Figure 2h**). This increase, however, was similar at genes regulated by TIP5 in ESC+2i and at random genes (**Supplementary Figure 2i**).

These results show that TIP5 specifically affects gene expression of ESC+2i and this process does not depend on the low DNA methylation content and low H3K27me3 levels at PRC targets characterising ground-state ESCs.

### TIP5 associates with large and active genomic regions of ESCs

To determine how TIP5 specifically regulates gene expression in ground-state ESCs, we measured and compared TIP5 genomic occupancy between ESC+2i and ESC+serum. We established an ESC line containing a FLAG-HA (F/H) tag at the N-terminus of both *Tip5* alleles (**Supplementary Fig. 3a-c**). Previous studies showed that the fusion of the F/H peptide at the N-terminus of TIP5 does not affect its activity ^22, 26^. The obtained ESC lines (F/H-TIP5-ESC) express TIP5 at levels similar to wild-type (wt) cells (**Fig. 3a**). Furthermore, cell morphology, proliferation, and expression of pluripotency genes were similar in both F/H-TIP5-ESC and parental ESC (data not shown). We performed ChIPseq analysis with anti-FLAG or anti-HA immunoprecipitation and assessed the specificity of TIP5-ChIP by comparing FLAG- or HA-ChIP of F/H-TIP5-ESCs and wt-ESCs. ChIP results were consistent with previous data showing a lack of TIP5 association with rRNA genes in both ESC+2i and ESC+serum (**Fig. 3b**) ^27^. The genomic binding pattern of TIP5 in both ESC+2i and ESC+serum did not appear as a distinct peak-like profile typical of transcription factors, but was rather enriched over large domains that extended up to several hundred kb (average size 33 kb, **Fig. 3c**). In contrast to differentiated cells, where TIP5 associates with repressive epigenetic signatures (i.e. DNA methylation and K3K9me2/3) ^20^, we found that TIP5 occupies active regions of the genome marked by H3K4me1 and H3K27ac and its occupancy correlates poorly with H3K27me3 and H3K9me3 levels in ESC+2i (**Figures 3c,d**). Consistently, TIP5-bound regions were excluded from the repressed lamina-associated domains (LADs) ^28^ (**Supplementary Fig. 3d**). While genomic distribution was similar in ESC+serum and ESC+2i (>88% commonly bound region) (**Supplementary Fig. 3e**), the amount of TIP5 binding was higher in ESC+2i (**Fig. 3b,c,e**). We validated the levels of TIP5 binding at selected genomic regions of ESC+2i and ESC+serum by ChIP experiments followed by quantitative (q)PCR measurements (**Fig. 3b**). In total, we found 1263 TIP5-bound regions that were unique to either of the two ESC states (**Fig. 3b, Supplementary Fig. 3e**). Most of ESC state specific TIP5-bound regions were found in ESC+2i (1003 TIP5 2i-specific sites, 80% of all unique TIP5-bound regions). Lack of TIP5 binding at these regions in ESC+serum associates with a decrease in H3K27ac or an increase in H3K27me3 content compared to ESC+2i, indicating that the association of TIP5 with ESC chromatin might depend on an active chromatin signature (**Fig. 3f,g**). Collectively, these results indicate that TIP5 associates with large and active chromatin domains, which are characterized by high H3K27ac and low H3K27me3 contents. Furthermore, despite the specific dependency of ESC+2i on TIP5 for gene expression, proliferation and differentiation capacity, TIP5 genomic occupancy is similar in these two ESC types. To gain insights on how TIP5 binding impacts gene expression, we investigated the position of TIP5-bound sites relative to differentially expressed genes in ESC+2i. We found that TIP5-bound regions in ESC+2i were depleted in the vicinity of TIP5 2i-regulated genes (log_2_(fold)= −0.77, FDR 10^−4^) (**Fig. 3h**). Moreover, we did not observe any evident positive correlation for linear genomic distance between TIP5-bound enhancers and TIP5 2i-regulated genes (**Supplementary Fig. 3f,g**). These results are also consistent with the ChIPseq analyses showing that TIP5 2i-regulated genes were enriched for H3K27me3 and depleted in H3K27ac (**Fig. 3i,j**), a chromatin signature that anticorrelates with the active chromatin regions bound by TIP5 (**Fig. 3c,d,f**). We conclude that the gene expression changes associated with TIP5 in ESC+2i are not a consequence of the direct binding of TIP5 to cis-acting regulatory elements (promoters and enhancers) of TIP5-2i regulated genes.

**Figure 3.**
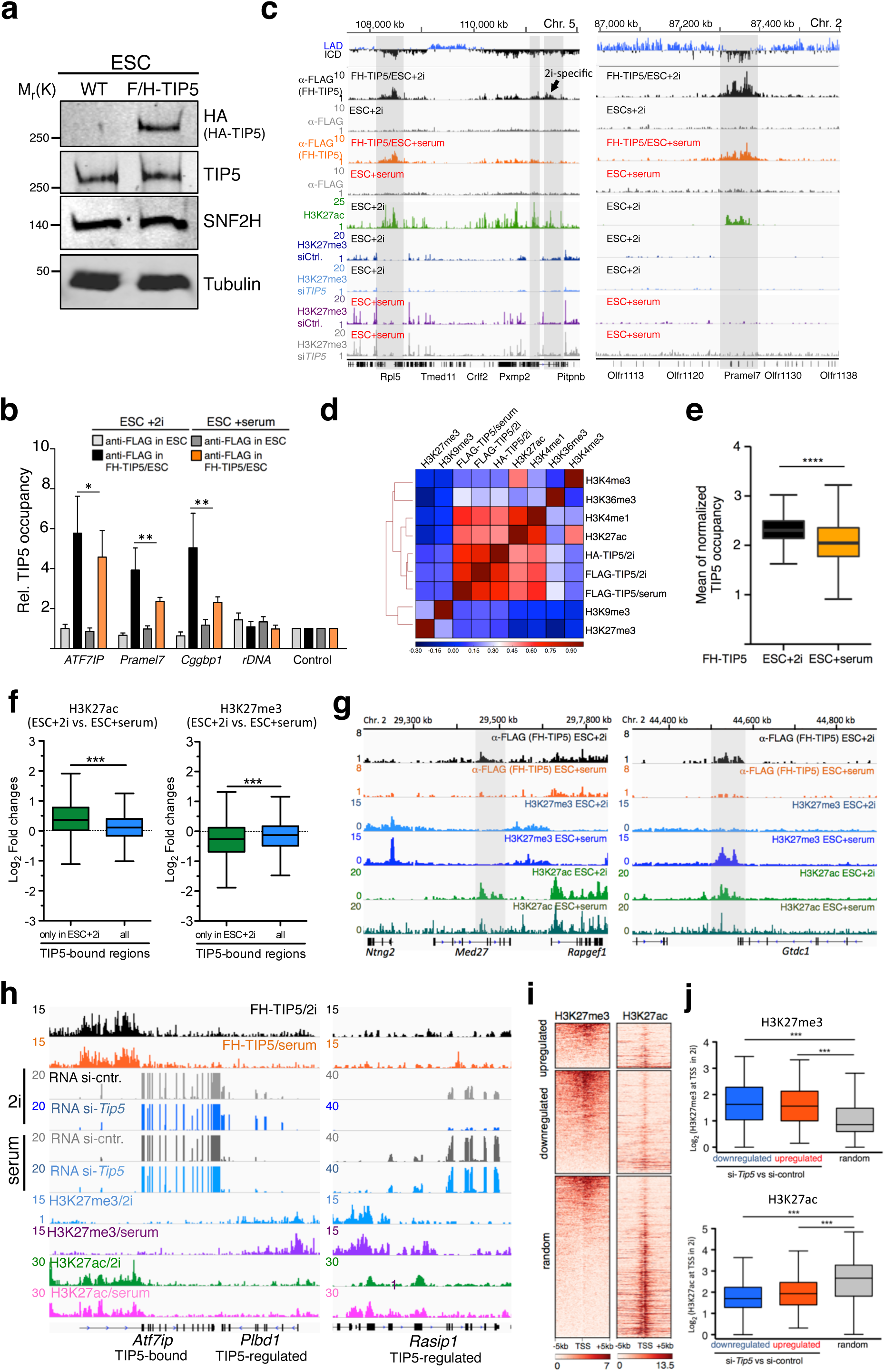
TIP5 associates with large and active chromatin domains in ESCs. **a.** Western blot showing equal TIP5 levels in wt-ESC and F/H-TIP5 ESC line containing FLAG-HA sequences at the N-terminus of both *Tip5* alleles. Whole cell lysates from equivalent amounts of cells were analysed. SNF2H and Tubulin are shown as protein loading controls. **b.** Anti-FLAG ChIP of wt-ESCs and F/H-TIP5-ESCs cultured in 2i or serum. Data were measured by qPCR and normalized to input and a control region that is not bound by TIP5. Average values of three independent experiments. Error bars represent s.d. and statistical significance (*P*-values) was calculated using the paired two-tailed t-test (*<0.05, **< 0.01). **c.** Representative images showing the association of TIP5 at regions enriched in H3K27ac and low in H3K27me3. The grey rectangles highlighted some of TIP5 associated regions. **d.** Pearson correlation heat map for the indicated ChIPseqs in ESC+2i. TIP5 ChIPseqs were performed with FLAG antibodies in *F/H-*TIP5-ESCs cultured in 2i or serum and with HA antibodies in *F/H-*TIP5-ESC+2i. Data of H3K4me3, H3K27ac and H3K27me3 are from this work. H3K36me3, H3K9me3 and H3K4me1 values in ESC+2i were taken from published data sets ^16, 17^. **e.** Boxplot showing the mean of normalized TIP5 occupancies in ESC+2i and ESC+serum at TIP5-bound regions. Statistical significance (*P*-values) for the experiments was calculated using the unpaired two-tailed t-test (**** < 0.0001). **f.** Differential TIP5-binding in 2i and serum correlates with H3K27ac occupancies. Boxplot showing the mean of normalized H3K27ac and H3K27me3 read counts in ESC+2i and ESC+serum at all TIP5-bound regions and TIP5-bound regions only in 2i were calculated and their log2-fold changes are plotted. H3K27ac in ESC+serum were taken from ^17^. Statistical significance (*P*-values) was calculated using the paired two-tailed t-test (***< 0.001). **g.** Representative images showing 2i-specific TIP5-bound regions and H3K27ac and H3K27me3 in ESC+2i and ESC+serum. **h.** Representative images showing that genes regulated by TIP5 are enriched in H3K27me3, depleted of H3K27ac and not bound by TIP5. **i.** Heat map profiles of H3K27me3 and H3K27ac at ±5 kb from the TSS of genes differentially regulated by TIP5 in ESC+2i and random genes. Differentially regulated genes are shown as upregulated or downregulated in ESC+2i upon TIP5 knockdown. Data were ranked by H3K27me3 levels. Transcription start site (TSS). **j.** Boxplot showing the mean of normalized H3K27me3 and H3K27ac read counts over +/− 1kb of TSSs from upregulated, downregulated and random genes.

### TIP5 bound regions mark the strongest long-range contacts in ESCs

Since TIP5 binds large active chromatin domains that are distal to the cis-regulatory elements of genes regulated by TIP5 in ESC+2i, we tested whether TIP5 plays a role in the genome organization of ground-state ESCs. We investigated whether TIP5-bound regions in ESCs display a preferential spatial genome compartmentalization. First, we made use of recently published high resolution (<750 bp) HiC map in ESC+serum, the highest to date in mammalian cells ^8^. Consistent with TIP5 binding to regions enriched in active histone modifications, we found that 96% of TIP5-bound regions are found in the active A compartment (**Fig. 4a**). Strikingly, we found stronger far-*cis* contacts between TIP5-bound regions in the genome, in general, and within A compartment, in particular, compared to near-*cis* contacts and interactions of regions not bound by TIP5 genome wide (**Fig. 4b,c**). These long-range contacts marked by TIP5 binding are specific to ESCs as they were gradually lost during the progression of neural differentiation (neural progenitors, NPCs, and cortical neurons, CNs) (**Fig. 4c,d Supplementary Figures 4a**).

**Figure 4.**
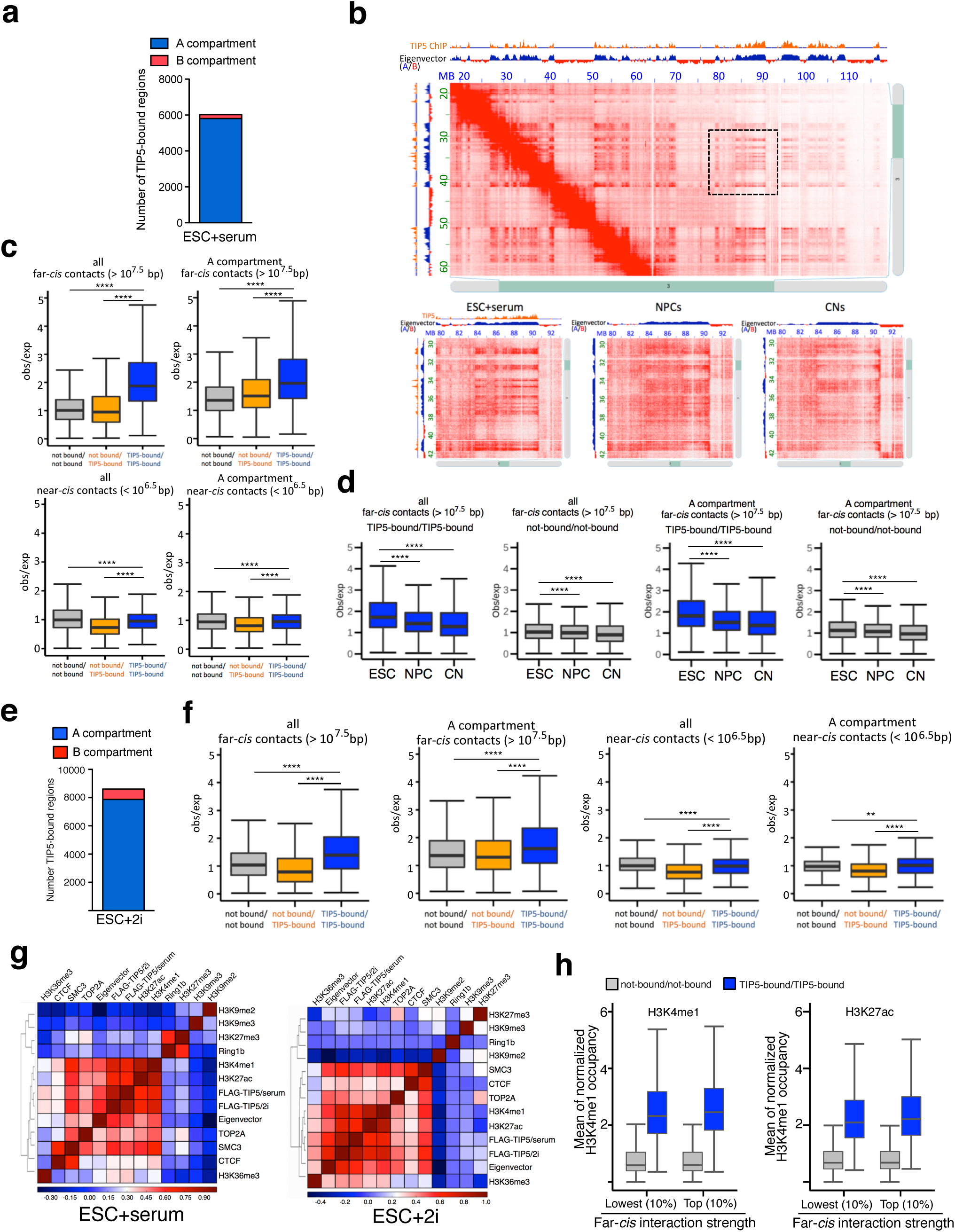
TIP5-bound regions mark the strongest long-distance contacts in ESCs. **a.** Number of TIP5-bound regions in A and B compartments of ESC+serum. **b.** Hi-C contact matrices for a zoomed in region on chromosome 6 at 100-kb resolution showing the presence of strong long-distance contacts corresponding to regions in A compartment bound by TIP5 in ESC+serum. Low panels. HiC contacts of a zoomed region at 50-kb resolution in ESC+serum, NPCs and CNs are shown. Data are from ^8^. HiC contacts were visualized using Juicebox ^2^. **c.** TIP5-bound regions mark A subcompartments that are characterized by the strongest far-*cis* contacts in ESC+serum. Boxplots indicating observed/expected contact values between TIP5-bound regions and loci not bound by TIP5 at near-*cis* and far-*cis* contacts in all genome or in A compartments. Statistical significance (*P*-values) was calculated using the unpaired two-tailed t-test (****< 0.0001). **d.** Long-range contacts marked by TIP5 binding are specific for ESCs. Boxplots indicating observed/expected contact values between TIP5-bound regions and loci not bound by TIP5 at far-*cis* contacts in all genome or in A compartments of ESC+serum, NPC and CN. Statistical significance (*P*-values) was calculated using the unpaired two-tailed t-test (****< 0.0001). **e.** Number of TIP5-bound regions in A and B compartments in ESC+2i. **f.** TIP5-bound regions mark A subcompartments that are characterized by the strongest far-*cis* contacts in ESC+2i. Boxplots indicating observed/expected contact values between TIP5-bound regions and loci not bound by TIP5 at near-*cis* and far-*cis* contacts in all genome or in A compartments of ESC+2i. Statistical significance (*P*-values) was calculated using the unpaired two-tailed t-test (** <0.001, ****< 0.0001). **g.** TIP5 binding correlates the best with eigenvector values. Heat map showing Pearson correlation values of ChIPseq data sets with the eigenvector values from HiC data in ESC+serum ^8^ and ESC+2i, respectively. **h.** Boxplots showing H3K4me1 and H3K27ac content of TIP5-bound regions and loci not bound by TIP5 at far-*cis* contacts in ESC+2i. Data from regions with lowest 10% and top 10% contact strength are shown.

Given the differences between TIP5 function in ESC+serum and ESC+2i, we next performed HiC analysis of ESC+2i (three independent biological experiments, 95 million pair end reads). Despite the relative lower resolution of these HiC maps relative to the HiC data of ESC+serum ^8^, the conclusions of the analysis of both datasets were similar. A and B compartments annotated in ESC+2i (our HiC) and in ESC+serum (HiC from ^8^) were well correlated, indicating that despite the differences in chromatin states between the two ESC types genome compartmentalization is similar (**Supplementary Fig. 4b**). Moreover, ESC+2i and ESC+serum showed a similar contact profile, including higher far-*cis* contacts at A compartments compared to near-*cis* contacts, which is consistent with previous results showing the propensity of active chromatin of ESCs to segregate and establish long-range interactions ^8^ (**Supplementary Fig. 4c**). We found that TIP5-bound regions in ESC+2i are also found within A compartments (**Fig. 4e**). Consistent with our analysis in ESC+serum, also in ESC+2i far-*cis* contacts between TIP5-bound regions are the strongest relative to near-*cis* contacts and that between regions not bound by TIP5 (**Fig. 4f, Supplementary Fig. 4d**). Therefore, although chromatin is more open and epigenetically active in ESC+2i than in ESC+serum, genome compartmentalization and TIP5-bound far-*cis* contacts are similar in both ESC types.

TIP5-bound regions within the A compartment of ESC+2i and ESC+serum had higher eigenvector values than regions not bound by TIP5 (**Supplementary Fig. 4e**). Remarkably, the correlation between ChIPseq signal enrichment for TIP5 binding and histone marks and eigenvector value indicated that TIP5 has the strongest correlation with positive eigenvector values in both ESC+2i and ESC+serum (**Fig. 4g**). The presence of A and B subcompartments with distinct patterns of histone modifications has been described in human differentiated cells ^2^. As TIP5 associates with H3K27ac and H3K4me1 (**Fig. 3**), these A subcompartments represent regions particularly enriched in active histone marks. Strikingly, however, the strength in far-*cis* contacts between TIP5-bound regions and regions not bound by TIP5 does not depend on H3K27ac and H3K4me1 levels (**Fig. 4h**). Taken together, the results indicate that TIP5 binds active regions of A compartments that have the strongest long-range interactions in both ESC types.

### TIP5 counteracts the propensity of active chromatin of ground-state ESCs to invade repressive domains

The binding of TIP5 within active chromatin regions engaged in long-range contacts in ESCs, suggested to us that TIP5 binding within active chromatin domains, in ground-state but not developmentally advanced ESCs, could impact chromatin states and contribute to the gene expression changes observed specifically in ESC+2i. To test this, we first measured chromatin accessibility upon TIP5 knockdown in both ESC types using ATACseq. As expected, the most accessible chromatin regions in both ESC types were found in A compartments (72% in ESC+2i, >75% in ESC+serum). About one third of accessible chromatin sites coincided with TIP5 bound regions, supporting the previous analyses showing that TIP5 associates with active chromatin regions that are known to have elevated chromatin accessibility (**Fig. 5a, Fig. 3, Supplementary Fig. 5a,b**). ESC+2i displayed higher chromatin accessibility in B compartment compared to ESC+serum, underscoring the more open chromatin state of ESC+2i even at repressive domains (**Fig. 5b**). Increased chromatin accessibility correlates with elevated H3K27ac levels in both ESC types, indicating that this modification specifically marks distinct open chromatin domains in the two pluripotency states (**Fig. 5c,d**). Remarkably, depletion of TIP5 induced changes in chromatin accessibility of ESC+2i whereas ESC+serum were unaffected, indicating that TIP5 specifically modulates chromatin structure in ground-state ESCs (**Fig. 5e,f, Supplementary Fig. 5c**). Overall TIP5 knockdown increased chromatin accessibility in ESC+2i (1952 increased chromatin accessibility sites vs. 440 decreased accessibility sites) (**Fig. 5e,f**). Decreased accessibility sites (DAS) were close to genes downregulated upon TIP5 knockdown, indicating reduced chromatin accessibility at genes that undergo transcriptional repression in the absence of TIP5 whereas no correlation was found in increased accessibility sites (IAS) (**Fig. 5g**). A large portion of DAS (29.28%) was located at gene promoters whereas only 2.94% of IAS were present at these genomic elements (**Fig. 5h**). Almost half (44%) of IAS were located within the B compartment, which is not bound by TIP5 (**Fig. 5e**). We did not find enrichment in IAS or DAS at TIP5 bound regions, indicating that changes in chromatin accessibility upon TIP5-KD are not dependent on the direct interaction of TIP5 with these regions (**Supplementary Fig. 5d**). Remarkably, the position of IAS was not homogenously distributed within the compartments (**Fig. 5i, Supplementary Table 3**). In B compartments, IAS were only enriched at B boundaries whereas DAS did not show any specific positional enrichment in the A compartment (**Fig. 5i, Supplementary Table 3**). Furthermore, we found that also genes upregulated upon TIP5 depletion in ESC+2i were enriched in B compartment relative to all expressed genes and were preferentially located at B boundaries (**Fig. 5i,j, Supplementary Table 3**). Consistent with the increased chromatin accessibility and upregulation of gene expression in B compartment upon TIP5 depletion, we found that H3K27ac and H3K4me3 levels also increase in B compartments (**Fig. 5k**).

**Figure 5.**
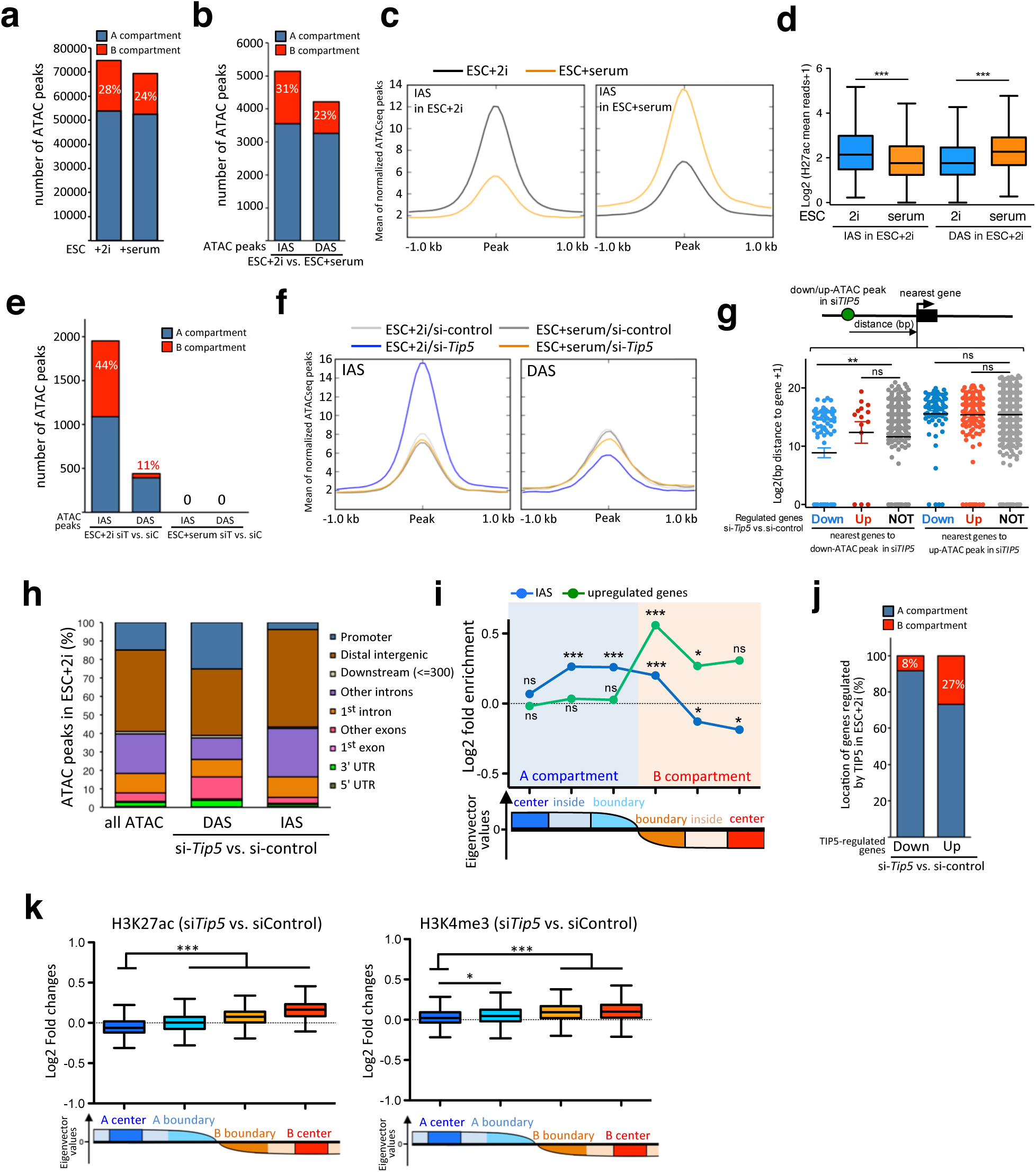
TIP5 regulates chromatin accessibility in ground-state ESCs. **a.** Numbers of ATAC peaks called separately in ESC+2i and ESC+serum including their localization according to compartmentalization. **b.** Numbers of significant changed ATAC peaks in ESC+2i vs. ESC+serum and their localization according to compartmentalization. Increased chromatin accessibility sites (IAS), decreased accessibility sites (DAS). **c.** Average profile of normalized ATAC read counts over differential peaks in ESC+2i and ESC+serum changed ATAC peaks. The signal is plotted over +/− 1kb on the center of the ATAC peak. **d.** H3K27ac correlates with differential accessible sites between ESC+2i and ESC+serum. Boxplot showing the mean values of normalized H3K27ac at ATAC peaks (+/− 500bp) that are increased and decreased in ESC+2i compared to ESC+serum, respectively. Statistical significance (*P*-values) for the experiments was calculated using the paired two-tailed t-test (***<0.001). **e.** TIP5 depletion increases chromatin accessibility only in ESC+2i. Numbers of significant changed ATAC peaks in ESC+2i and ESC+serum upon TIP5 knockdown and their localization according to compartmentalization. **f.** Average density plots of normalized ATAC read counts over differential ATAC peaks of ESC+2i treated with siRNA-*Tip5* and siRNA-control over changed ATAC peaks. The signal is plotted over +/− 1kb centered on the ATAC peak. **g.** The distances of increased (IAS) and decreased (DAS) chromatin accessibility sites in TIP5-depleted ESC+2i to the nearest gene promoter (+/− 5kb of TSS) were calculated. Distances in bp to the nearest promoter of genes upregulated, downregulated or not regulated upon TIP5 knockdown in ESC+2i are plotted. Statistical significance (*P*-values) for the experiments was calculated using the unpaired two-tailed t-test (*** < 0.001). **h.** Genomic annotation of IAS and DAS in ESC+2i depleted of TIP5 perfomed with ChIPseeker ^58^. **i.** Fold-enrichment relative to expected of regions with increased chromatin accessibility (IAS) and upregulated genes, upon TIP5 KD, at sub-regions within compartment A and B (center, inside and boundaries). Compartments smaller than 500 kb were excluded from the analysis. **j.** TIP5-upregulated genes are enriched in the B compartment. Numbers of genes upregulated and downregulated upon TIP5 knockdown in ESC+2i including their localization according to compartmentalization. **k.** Active histone marks increase in the B compartment upon TIP5 knockdown. Boxplots indicating the log2 fold-changes of normalized H3K27ac and H3K4me3 read counts in ESC+2i treated with siRNA-*Tip5* and siRNA-control at boundaries and centers of A and B compartments.

Taken together these results indicated that TIP5 regulates chromatin accessibility specifically in ESC+2i. Specifically, our data highlighted the role of TIP5 in limiting invasion of active chromatin into the repressive chromatin of B compartment in ESC+2i.

### TIP5 is required for 3D genome architecture of ground-state ESCs

To determine whether the role of TIP5 in controlling active chromatin domains was a result of its function in genome organization of ground-state ESCs, we performed HiC analysis of ESC+2i in control and TIP5-depleted ESC+2i. For each group we performed three independent biological experiments that showed high similarity (**Supplementary Fig. 6a, Supplementary Table 4**). Analysis of ESC+2i depleted of TIP5 revealed a genome wide increase in far-*cis* contacts (>10Mb) compared to control cells (**Fig. 6a,b, Supplementary Fig. 6b**). Although these extremely long-range interactions occurred at a relatively low frequency, their increase upon TIP5-KD is consistent across all three independent HiC experiments, suggesting that the lack of TIP5 induced the formation of spurious genomic contacts. Increase in long-distance contacts could be observed in A and B compartments and at TIP5-bound regions and regions not bound by TIP5 (**Supplementary Fig. 6c**), a result that is consistent with the expansion of active chromatin, which in ESCs have been shown to be prone to form long-range contacts ^8^. Alterations in genome architecture mediated by TIP5 in ESC+2i were also evident by a decrease in absolute eigenvector values at the boundaries of B compartments whereas the center regions of B compartment and the entire A compartment were not significantly affected (**Fig. 6c**). These results are consistent with the increased chromatin accessibility and euchromatic marks observed particularly at B boundaries upon TIP5 depletion (**Fig. 5i,k**). Collectively, the results further support a role of TIP5 in protecting repressive compartments by the invasion of active domains.

**Figure 6.**
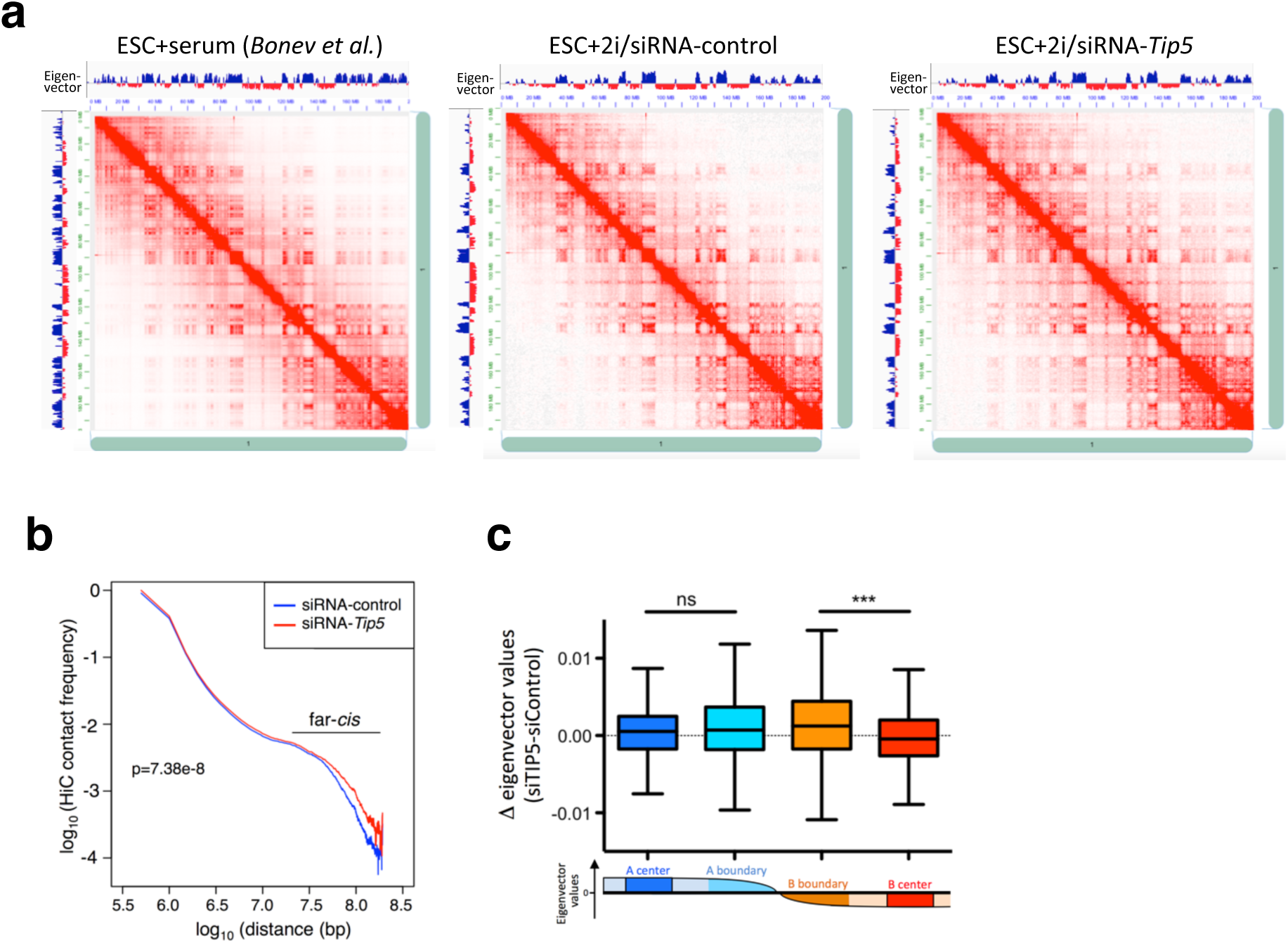
TIP5 limits spurious long-distant contacts in ground-state ESCs. **a.** Hi-C contact matrices for chromosome 1 of ESC+serum, ESC+2i/siRNA-control and ESC+2i/siRNA-*Tip5* showing the presence of strong long-distance contacts corresponding to regions in A compartment bound by TIP5 in ESC+serum. Eigenvector values are shown. **b.** Relative contact frequencies over genomic distances in ESCs treated with siRNA-control and siRNA-TIP5. **c.** Boxplot indicating differences in eigenvector values at boundaries and centers of A and B compartments. Differences in eigenvector values were calculated from HiC data sets from ESC+2i treated with siRNA-*Tip5* and siRNA-control.

### TIP5 regulates H3K27me3 occupancy in ground-state ESCs

To investigate whether TIP5 might impact the epigenetic landscape at repressed chromatin regions in ESC+2i, we analyzed H3K27me3 that is enriched at Polycomb domains. These regions are often flanked by CTCF boundaries, however, insulation mediated by CTCF does not act as a direct barrier to H3K27me3 spreading ^3, 29^. Western blot analyses indicated that total levels of H3K27me3 are not altered in the absence of TIP5 (**Fig. 7a, Supplementary 7a**). Strikingly, ChIPseq experiments revealed a global re-distribution of H3K27me3 in ESC+2i whereas ESC+serum were not affected (**Fig. 7b,c**). For promoters with high levels of H3K27me3, TIP5 depletion resulted in a decrease in H3K27me3, while promoters with relatively low H3K27me3 levels displayed an increase of this repressive mark (**Fig. 7b,d**). Quantitative H3K27me3 ChIPseq using *Drosophila* spike-in chromatin and ChIP-qPCR experiments validated the decrease of H3K27me3 at TSS of selected genes with high H3K27me3 content and showed that this effect was specific for ESC+2i (**Fig. 7e, Supplementary Fig. 7b**). Similar results were also obtained with a different siRNA-*Tip5* sequence (**Supplementary Fig. 7c**). Importantly, the redistribution of H3K27me3 did not correlate with transcriptional changes observed upon TIP5 knockdown as both upregulated and downregulated genes showed decreased H3K27me3 occupancies, a result that is consistent with the elevated H3K27me3 content at TSS of TIP5 2i-regulated genes (**Fig. 7f**). Furthermore, the changes in H3K27me3 could not account for the transcriptional changes since TIP5 depletion in *Ring1b*^−/−^ ESCs and *Eed*^−/−^ ESCs cultured in 2i still affected cell proliferation and expression of TIP5-2i regulated genes (**Supplementary Fig. 7d,e**). These results suggest that H3K27me3 alterations in TIP5 depleted ESC+2i could not be caused by a response to transcriptional differences. Accordingly, alterations in H3K27me3 occupancy was not restricted to TSSs but was also detected at intergenic regions as in the case of a subset of regions where CTCF sites mark transitions in H3K27me3 enrichment (**Fig. 7g**). Upon TIP5 depletion, H3K27me3 redistributed over CTCF sites, decreasing at high H3K27me3 regions and increasing at adjacent low H3K27me3 regions. Consistent with the specific role of TIP5 in ESC+2i, H3K27me3 transitions marked by CTCF sites in ESC+serum were not affected upon TIP5 knockdown. All these results were validated with independent H3K27me3 ChIP-qPCR experiments and another siRNA-*Tip5* sequence (**Fig. 7h, Supplementary Fig. 7c**). Given that the disruption of boundaries by CTCF depletion was shown not to affect H3K27me3 genome-wide occupancy ^3, 6^, the results suggest that insulation and reinforcement of H3K27me3 domains might occur through changes in genome organization mediated by TIP5 loss. To test this, we analysed the H3K27me3 domain *HoxA* gene cluster that is not bound by TIP5 and shows strong internal contacts in ESC+2i, forming a small sub-compartment that separates two larger TADs (**Fig. 7i**). Upon TIP5 depletion, this domain decreased H3K27me3 content and display alterations in contacts. Noteworthy, new contacts formed between the *HoxA* cluster and active domains previously bound by TIP5 and *HoxA1* and *HoxA7* genes became upregulated (**Fig. 2b, Supplementary Table 1, 5**). Structural changes were also observed at another H3K27me3 domain, the *HoxB* locus, showing loss of loop anchor points and fusion with the adjacent compartment (**Supplementary Fig. 7e**). Together, these results indicate that TIP5 regulates H3K27me3 occupancy specifically in ESC+2i. Alterations in H3K27me3 are not a direct consequence of transcriptional changes or TIP5 binding with these regions but instead are likely a consequence of the lack of TIP5 binding within active chromatin domains that affects genome organization.

**Figure 7.**
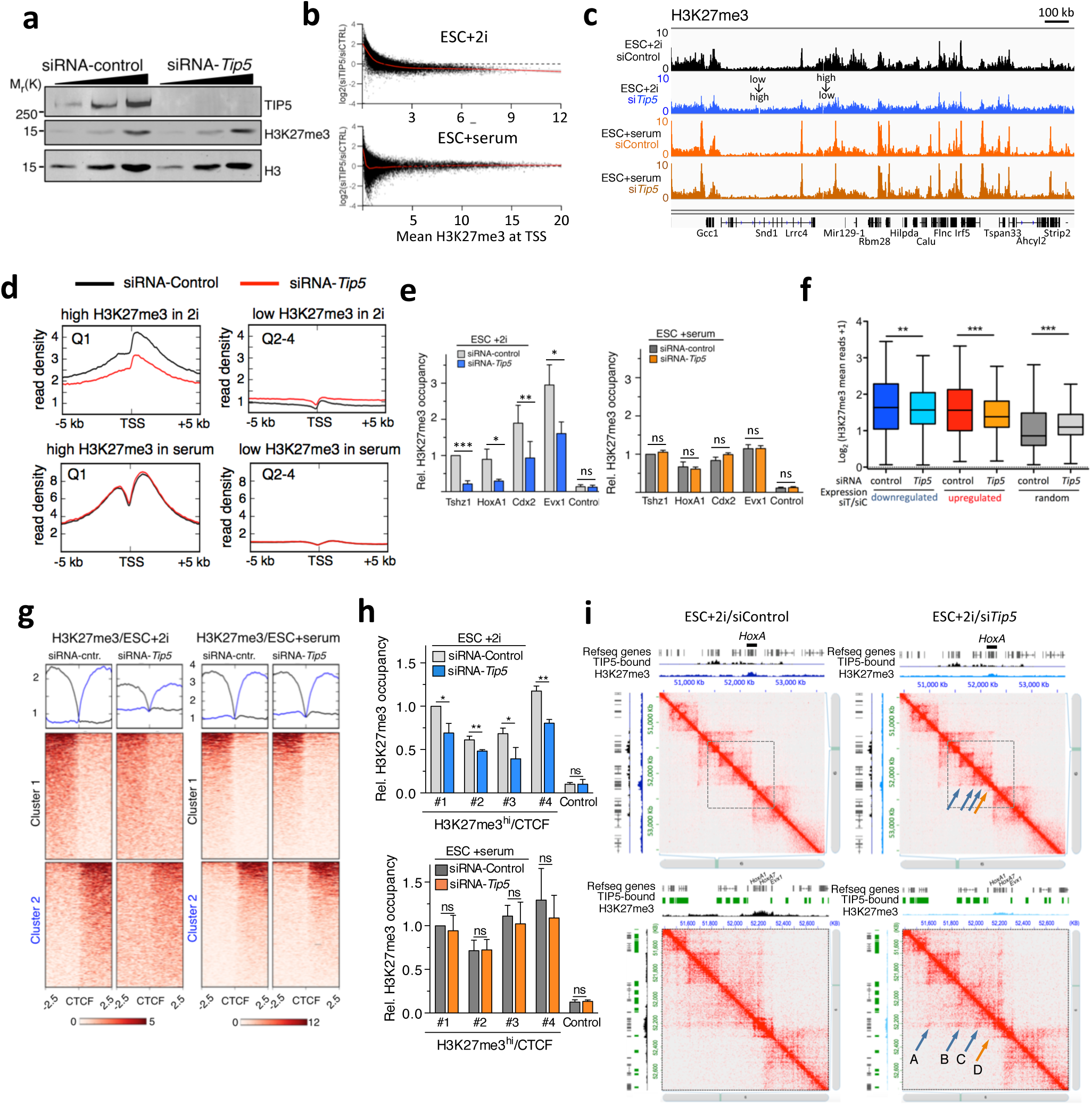
TIP5 regulates H3K27me3 occupancy in ground-state ESCs. **a.** Western blot showing total levels of H3K27me3 in ESC+2i transfected with siRNA-control and siRNA-*Tip5*. Histone H3 is shown as a protein loading control. **b.** Scatterplot showing the changes in H3K27me3 upon TIP5 depletion in ESC+2i and ESC+serum. Mean of normalized H3K27me3 read counts in ESC+2i and ESC+serum treated with siRNA-control or siRNA-*Tip5* over all TSSs (+/− 1kb) were calculated. Log2-fold changes of (siRNA-*Tip5/*siRNA-control) are plotted relative to the mean of normalized H3K27me3 occupancies. **c.** Representative images showing the H3K27me3 occupancy. **d.** Average density plot of ChIPseq read counts of H3K27me3 at −/+ 5 kb from the TSS of refseq genes in ESCs treated with siRNA-control or siRNA-*Tip5*. Q1: first quartile corresponding to sequences with high H3K27me3 content. Q2-4: quartiles 2-4 corresponding to sequences with low H3K27me3 content. **e.** ChIP-qPCR of H3K27me3 in ESC+2i and ESC+serum treated with siRNA-control or siRNA-*Tip5.* Data were measured by qPCR and normalized to input and to *Tshz1* value in control cells. Control represents an intergenic sequence that does not contain H3K27me3. Average values of three independent experiments. Error bars represent s.d. Statistical significance (*P*-values) for the experiments was calculated using the paired two-tailed t-test (ns: non significant, *<0.05, **< 0.01, ***<0.001). **f.** Boxplot showing changes in H3K27me3 in ESC+2i upon TIP5 at TIP5-regulated and random genes. Mean of normalized H3K27me3 read counts in ESC+2i treated with siRNA-control or siRNA-*Tip5* over TSSs (+/− 1kb) of upregulated, downregulated and random genes were calculated. Log2-fold changes of (siRNA-*Tip5/*siRNA-control) are plotted. Statistical significance (*P*-values) for the experiments was calculated using the paired two-tailed t-test (** < 0.01 and *** < 0.001). **g.** Average density and heatmap profile of H3K27me3 transition state regions marked by CTCF sites in ESC+2i or ESC+serum transfected with siRNA-control or siRNA-*Tip5*. Data were ranked by H3K27me3 content in the corresponding siRNA-control ESCs. Cluster 1 represents high to low H327me3 transition whereas Cluster 2 shows low to high H3K27me3 levels. **h.** H3K27me3 ChIP-qPCR in ESC+2i (top panel) or ESC+serum (bottom panel) transfected with siRNA-control or siRNA-*Tip5*. Values show H3K27me3 occupancy at four sequences (#1-#4) with elevated H3K27me3 at CTCF boundary (H3K27me3^hi^/CTCF). Data were measured by qPCR and normalized to input and to sequence #1 value in control cells. Average values of three independent experiments. Error bars represent s.d. Statistical significance (*P*-values) for the experiments was calculated using the paired two-tailed t-test (ns: non significant, *<0.05, **< 0.01). **i.** Hi-C contact matrices for a zoomed in region on chromosome 6 containing *HoxA* cluster (10-kb resolution in upper panels, 5 kb resolution in lower panels). Blue arrows indicated gain or increase of contacts in ESC+siRNA-*Tip5* compared to control cells (siRNA-control). Orange arrows indicate loss or decrease in contacts. Quantifications are shown in **Supplementary Table 5**.

### TOP2A interacts with TIP5 and its activity is required for TIP5 function

In order to understand how TIP5 affects genome organization and chromatin features in ESC+2i, we analysed which proteins interact with TIP5. Due to its tight association with chromatin, we aimed to purify TIP5 and its direct interaction partners in their native environment by establishing a protocol that allows immunoprecipitation from purified chromatin (Chromatin-IP, **Fig. 8A**). F/H-TIP5 and wt-ESC nuclei were incubated with the reversible protein-protein specific crosslinker dithiobis(succinimidyl propionate) (DSP) and chromatin was isolated by centrifugation of nuclear extracts. After digestion with MNase, solubilisation of mononucleosomes was achieved with 1% SDS, which does not affect protein-protein interactions stabilized by DSP crosslinking. The identification of TIP5-interacting proteins on chromatin was determined by comparing anti-FLAG immunoprecipitates from F/H-TIP5 and wt-ESC chromatin followed by mass spectrometry (**Fig. 8B, Supplementary 5**). Analysis from three independent experiments revealed 24 proteins that consistently interact with TIP5 in ESC+2i. As expected, the strongest interacting partner of TIP5 was SNF2H (**Fig. 8B**), suggesting that the function of TIP5 in ESCs can be directed via the remodeling complex NoRC complex. Moreover, we detected all known and previously validated TIP5-interactors in differentiated cells such as DNMT1, PARP1 and DHX9 ^20, 21, 25, 26^. Remarkably, among the top TIP5-interacting proteins we found Topoisomerase 2A (TOP2A) and structural maintenance of chromosomes protein 3 (SMC3), which is part of the cohesin complex (**Fig. 8B,C**). Furthermore, two out of three chromatin IP experiments detected the interaction of TIP5 with another cohesin component, SMC1 (**Supplementary Table 6**). Importantly, in the correlation analysis between ChIPseq signal enrichment and eigenvector values in ESCs, TOP2A and cohesin cluster together and were close to TIP5 and eigenvector values (**Fig. 3g**). Both TOP2A and cohesin are well known regulators of chromatin architecture. Topoisomerases exert the key function in relieving torsional stress of DNA ^30^. In ESCs, TOP2A is the most highly expressed type II isoenzyme and its inactivation is embryonically lethal ^31, 32^. We did not detect any interaction of TIP5 with CTCF, a result that is consistent with the lack of TIP5 enrichment at TAD boundaries and CTCF sites (data not shown). Interestingly, recent results revealed that abolishing cohesin loading to DNA resulted in an increase of far-*cis* chromatin contacts as we observed upon TIP5 depletion in ESC+2i ^5^.

**Figure 8.**
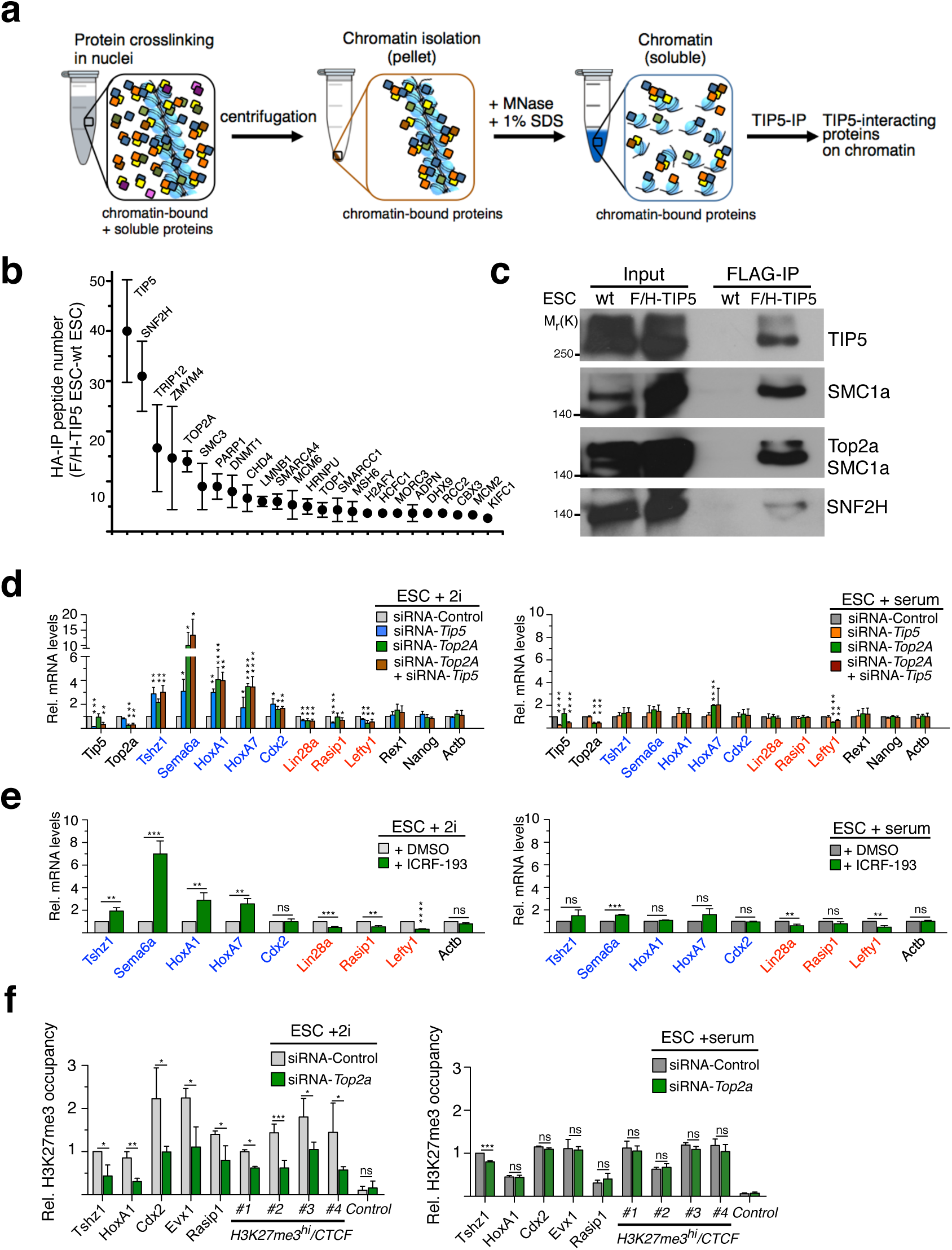
TIP5 associates with TOP2A and components of the cohesin complex on chromatin. **a.** Scheme representing the strategy used to identify protein interacting with TIP5 on chromatin (Chromatin-IP). **b.** TIP5-interacting proteins on chromatin. Mass spectrometry analysis of FLAG immunoprecipitates from wt and F/H-TIP5 ESC+2i. Values of peptide number are shown as the difference between peptides obtained in FLAG-IP of F/H-TIP5 and wt ESC chromatin extracts. Data are from three independent experiments. The proteins found enriched in FLAG-IP of F/H-TIP5 ESCs in all three experiments are shown. Further TIP5-interacting proteins identified in only two or one of the immunoprecipitation (IP) experiments are listed in **Supplementary Table 6**. **c.** Anti-FLAG IP of chromatin purified from wt- and F/H-TIP5-ESCs. Western blot shows the interaction of TIP5 with SNF2H, TOP2A and SMC1a. After protein transfer, membrane was first incubated with anti-SMC1a antibodies and after signal detection re-incubated with anti-TOP2A antibodies. **d.** TOP2A and TIP5 regulate gene expression in ESC+2i via a shared pathway. qRT-PCR of genes regulated by TIP5 in ESC+2i (left panel) and in ESC+serum (right panel) upon TOP2A or TIP5 knockdown by siRNA. Upregulated genes in ESC+2i upon TIP5-KD are labeled in blue, downregulated genes are in red. *Nanog*, *Rex1* and *Actin* B (*ActB*) are shown as genes not regulated by TIP5. mRNA levels were normalized to *Rps12* mRNA and to ESCs transfected with siRNA-Control. Average values of three independent experiments. Error bars represent s.d. Statistical significance (*P*-values) for the experiments was calculated using the paired two-tailed t-test (ns: non significant, *<0.05, **< 0.01, ***<0.001, ****<0.0001). **e.** Inhibition of type 2 topoisomerase activity phenocopies the alterations in gene expression of TIP5-regulated genes in ESC+2i observed upon TOP2A or TIP5 knockdown. ESCs were treated with the TOP2 inhibitor ICF-193. mRNA levels were normalized to *Rps12* mRNA and to ESCs treated with DMSO. Average values of three independent experiments. Error bars represent s.d. Statistical significance (*P*-values) for the experiments was calculated using the paired two-tailed t-test (ns: non significant, *<0.05, **< 0.01, ***<0.001, ****<0.0001). **f.** ChIP analysis showing that TOP2A regulates H3K27me3 occupancy in ESC+2i but not in ESC+serum. Data were measured by qPCR and normalized to input and to *Tshz1* value in cells transfected with siRNA-control. Control represents an intergenic region that does not contain H3K27me3. Average values of three independent experiments. Error bars represent s.d. Statistical significance (*P*-values) for the experiments was calculated using the paired two-tailed t-test (ns: non significant, *<0.05, **< 0.01, ***<0.001).

To determine whether TOP2A regulates ESC+2i similarly to TIP5, we analysed and compared gene expression in ESC+2i and ESC+serum upon knockdown of TOP2A or after treatment with ICRF-193, a potent TOP2 inhibitor ^33^. Depletion of TOP2A or its catalytic inhibition in ESC+2i induced transcriptional changes at candidate genes that mimicked those observed upon TIP5 knockdown (**Fig. 8d,e**). Moreover, the lack of apparent additive activation or repression of TIP5 2i-regulated genes upon combined depletion suggests that TIP5 regulates gene expression in ESC+2i with TOP2A via shared pathways. Importantly, depletion of TOP2A or treatment with ICRF-193 in ESC+serum had either a moderate effect (*HoxA1* and *Lefty1*) or no effect in gene expression of the analyzed TIP5 2i-regulated genes. Finally, H3K27me3 changes upon TOP2A depletion were similar to those observed upon TIP5 knockdown (**Fig. 8f**). Also in this case, alterations in H3K27me3 were observed only in ESC+2i but not in ESC+serum.

Depletion of cohesin component SMC3 in ESC+2i caused similar changes in gene expression of TIP5 2i-regulated genes (**Supplementary Figure 8a**). In ESC+serum, however, SMC3 depletion could still induce activation of *Sema6a* and only a moderate upregulation of *HoxA7*. In both ESCs depleted of SMC3, TIP5 levels were 30% reduced. Although the reduction in TIP5 levels might have an influence on the analysis of TIP5 2i-regulated genes by SMC3 depletion in ESC+2i, this decrease was not sufficient to affect H3K27me3 occupancy since upon SMC3 knockdown H3K27me3 level remain the same, a result that is consistent with previous studies ^6^ (**Supplementary Figure 8b**).

To determine how TOP2A and SMC3 act together with TIP5, we analyzed whether they are required for the association of TIP5 with chromatin. We performed TIP5-ChIP analyses in ESC+2i and ESC+serum and found that TOP2A depletion as well as inhibition of topoisomerase activity decreased the association of TIP5 with TIP5-bound sequences in both ESC+2i and ESC+serum (**Supplementary Figures 8c-e**). SMC3 depletion reduced the association of TIP5 with chromatin in ESC+2i whereas this effect was relatively modest in ESC+serum, further highlighting the difference between the chromatin of these two pluripotency state. In contrast, TIP5 depletion did not affect the occupancy of SMC1a at selected cohesin bound regions in ESC+2i (**Supplementary Figures 8f**). The same analysis of TOP2A occupancy could not be performed due to technical issues in generating a specific TOP2A ChIP signal.

The association of TIP5 with two important chromatin regulators such as TOP2A and cohesin provides further evidence that TIP5 acts on chromatin to regulate genome organization in ground-state ESCs. Furthermore, the data show that TOP2A activity is required for TIP5 binding to chromatin thereby phenocopying the two readouts of TIP5 function in ESC+2i, regulation of gene expression and H3K27me3 occupancy.

## Discussion

Chromatin has the intrinsic property to segregate into subnuclear compartments forming long-distance contacts between domains with similar chromatin states ^2^. In this work, we made use of the two closely and developmentally related ESC types, ground-state and developmentally advanced ESCs, to elucidate how their genome organization is regulated. We showed that both pluripotent ESC types display similar genome compartmentalization despite their different chromatin and epigenetic features However, only ground-state ESCs rely on TIP5, a component of the nucleolar remodeling complex NoRC^20, 21^. The role of TIP5 appeared ground state-specific since the absence of TIP5 in ESC+2i, but not in ESC+serum, led to changes in cell proliferation and differentiation, chromatin structures, H3K27me3 occupancy and gene expression. Importantly, alterations in chromatin accessibility, H3K27me3 occupancy and gene expression in ESC+2i are not a consequence of binding of TIP5 to any of these affected sites. One example of this is that TIP5 depletion in ESC+2i leads to increased chromatin accessibility and upregulation of genes at the repressive B compartments that are not bound by TIP5. We showed that TIP5 binds to large and active regions of A compartments that strongly intersect through long-range contacts in ESCs. Therefore, it appears that the function of TIP5 in ESC+2i is on active and highly interacting domains but the effects observed upon TIP5 depletion are mostly restricted to repressive chromatin regions. These results suggest a model where TIP5 bound to active chromatin domains counteracts their expansion into repressed domains and this function is required only in ground-state ESCs. The possibility that active chromatin can regulate heterochromatin through indirect mechanisms has also been recently proposed by showing that MRG-1, a factor exclusively bound to euchromatin, drives spatial sequestration of heterochromatin in *C. elegans* nuclei ^34^. Given the striking similarity in the TIP5 genomic occupancy in both pluripotent states, we propose that TIP5 function is determined by the state of its substrate (i.e. chromatin) rather than by TIP5 itself. Relative to ESC+2i, chromatin of ESC+serum contains a more repressive epigenetic state and more condensed chromatin structures at the level of nucleosomes ^12–16^. This structure might be sufficient to limit excessive chromatin flexibility, making ESC+serum less dependent on factors, such as TIP5, to limit the invasion of active domains into repressive regions.

Our data indicated that the tight control of genomic compartmentalization is central for the regulation of gene expression, epigenetic landscapes and stem cell integrity. In this regard, we observed that upon TIP5 depletion ESCs+2i proliferate significantly slower and had impaired differentiation potential, which is an important feature of pluripotency. Importantly, neither DNA methylation nor H3K27me3 have an influence on proliferation and transcription of genes regulated by TIP5 in ESC+2i. Furthermore, changes in H3K27me3 were not a consequence of transcriptional changes since the decrease of H3K27me3 occurred at the promoters of both up- and downregulated genes and at intergenic domains as in the case of a subset of regions where CTCF sites mark transitions in H3K27me3 enrichment. It is important here to note that previous studies indicated that CTCF does not act as a direct barrier to H3K27me3 spreading ^3, 29^. The decrease of H3K27me3 content and structural changes at the Polycomb domains *HoxA* and *HoxB* loci that are not bound by TIP5 suggests that insulation and reinforcement of H3K27me3 domains might occur through 3D genomic contacts. Accordingly, recent results showed that regulation of gene expression by Polycomb in ESCs also occur in a 3D interaction network ^18^.

How TIP5 acts on active chromatin domains in ground-state ESCs to limit their invasion into repressive domains remains elusive and will be the focus of future studies. However, our chromatin-IP analysis revealed that the strongest interacting partner of TIP5 was SNF2H, the only other component of the remodeling complex NoRC, suggesting that chromatin remodeling activities can play a role in this process. Furthermore, our results suggest that this mechanism most likely involves catalytically active TOP2A that interacts with TIP5 on chromatin and is required for TIP5 binding, thereby affecting gene expression and H3K27me3 occupancy similarly to TIP5. Interestingly, inhibition of TOP2 activity was shown to increase chromatin accessibility at H3K27me3 sites in ESCs, suggesting that TOP2 has a role in making facultative heterochromatin inaccessible genome-wide ^35^. Furthermore, recent computational modeling studies suggest that TOP2 activity could enable loop extrusion by resolving torsional stress, proposing an active role of TOP2 in restricting genome compartmentalization ^36, 37^. Therefore, the relief of chromatin torsional stress mediated by TOP2A might facilitate not only the access of TIP5 on active chromatin but also to protect repressive domains by the invasion of euchromatin.

Taken together, our results showed that open and active chromatin domains have the propensity to invade repressive regions. The involvement of TIP5, as part of NoRC complex, and TOP2A suggest that this process can be limited by chromatin remodeling and the relief of chromatin torsional stress in order to preserve active and repressed genome partitioning and cell function and identity.

## Materials and Methods

### Cell culture

129/OlaHsd mouse embryonic stem cells were cultured in either 2i media composed of DMEM-F12 and Neurobasal medium (1:1, Life Technologies), supplemented with 1× N2/B27 (Life Technologies), 1× penicillin/streptomycin/l-glutamine (Life Technologies), 50 µM β-mercaptoethanol (Life Technologies), recombinant leukemia inhibitory factor, LIF (Polygene, 1,000 U/ml) and MEK and GSK3β inhibitors, 2i (Sigma CHIR99021 and PD0325901, 3 and 1 µM, respectively) or in serum medium containing DMEM (Life Technologies), 15% FCS, 1× MEM NEAA, 100 µM β-mercaptoethanol, 1× penicillin/streptomycin (Life Technologies). ESCs were seeded at a density of 50,000 cells/cm^2^ in culture dishes (Corning^®^ CellBIND^®^ surface) coated with 0.1% gelatin without feeder layer. Propagation of cells was carried out every 2 days using enzymatic cell dissociation. When indicated, ESCs were treated with 1 μM Staurosporine 16 hours. 2i conversions were carried out for 8 days by plating the same number of ESC-wt and *Tip5*-KO ESCs every 2 or 3 days.

ESCs were differentiated by culturing for 48–72h in complete medium: DMEM, 10% FCS, 1 mM sodium pyruvate (Sigma), 1× NEAA (Life Technologies), 1× penicillin/streptomycin/l-glutamine, 100 µM β-mercaptoethanol on 0.1% gelatin-coated culture dishes.

Differentiation toward neural progenitor cells (NPCs) was obtained according to previously established protocol ^38^. In brief, differentiation employed a suspension-based embryoid bodies formation (Bacteriological Petri Dishes, Bio-one with vents, Greiner^®^) in neural differentiation media (DMEM, 10% FCS, 1× MEM NEAA, 1× penicillin/streptomycin/l-glutamine, 100 µM β-mercaptoethanol). During the 8-day differentiation procedure, media were exchanged every 2 days. In the last 4 days of differentiation, the media were supplemented with 2µM retinoic acid (RA) to generate neuronal precursors. DNMT-TKO ESCs were kindly provided by M. Okano ^39^. *Eed*-KO and *Ring1b*-KO ESC+2i were a gift from C. Ciaudo (ETH, Zurich).

### Transfections

ESCs were transfected with the indicated siRNAs (50 nM siRNA) using Lipofectamine^®^ RNAiMAX (Life Technologies) in Opti-MEM^®^ GlutaMAX™ (Life Technologies) reduced-serum medium. ESCs were seeded 2h prior to siRNA transfection and collected 3 days post transfection if not stated differently. For analysis of differentiation potential of TIP5-depleted ESCs, transfected cells were re-seeded at equal cell numbers into complete differentiation medium 48h post transfection. Survival of differentiated cells was assessed 72h later. Efficiencies of siRNA-mediated depletions were monitored by qRT–PCR 3 days post-transfection. To measure cell proliferation, same number of cells were plated and counted after 3 days of siRNA treatment.

### Alkaline phosphatase (AP) staining

Cells were fixed in 4% paraformaldehyde for 10 min, washed with AP buffer (100 mM Tris-Cl pH 9.5, 100 mM NaCl, 50 mM MgCl2), and then incubated for 5–10 min in BCIP^®^/NBT liquid substrate system (Sigma). The reaction was blocked with 10 mM Tris and 1 mM EDTA for 10 min.

### ICRF-193 treatment

ESCs were seeded 24h prior to treatment. ICRF-193 was added directly to the medium to final concentration of 500nM, as it has previously been reported for ESCs ^40^. Cells were harvested for respective experiments after 24h of inhibitor treatment.

### RNA extraction, reverse transcription, and quantitative PCR (RT–qPCR)

RNA was purified with TRIzol reagent (Life Technologies). 1µg total RNA was primed with random hexamers and reverse-transcribed into cDNA using MultiScribe™ Reverse Transcriptase (Life Technologies). Amplification of samples without reverse transcriptase assured absence of genomic or plasmid DNA (data not shown). The relative transcription levels were determined by normalization to *Rps12* or *beta-Actin* mRNA levels, as indicated. qRT–PCR was performed with KAPA SYBR^®^ FAST (Sigma) on a Rotor-Gene Q (Qiagen). Primer sequences are listed in **Supplementary Table 7**.

### Chromatin fractionation

ESCs were collected by trypsinization, washed once with PBS and counted. ES cell pellets were resuspended at a concentration of 10mio cells/ml in chromatin fractionation buffer (10mM Hepes pH 7.6, 150mM NaCl, 3mM MgCl_2_, 0.5% Triton X-100, 1mM DTT freshly supplemented with cOmplete^™^ Protease Inhibitor Cocktail (Roche)) and incubated for 30 minutes at room temperature rotating. Precipitated chromatin was fractionated by centrifugation. Total and chromatin fractionated samples were further processed by MNase (S7 Micrococcal nuclease, Roche) digest for ensuring sufficient genomic DNA fragmentation. All samples were incubated in 1x Laemmli buffer (10% glycerol, 10mM Tris pH 6.8, 2% SDS, 0.1mg/ml bromphenolblue, 2% β-mercaptoethanol) at 95°C for 5 minutes and were further analyzed by Western Blotting.

### Generation of FLAG/HA-TIP5 ES cell line by CRISPR/Cas9

CRISPR/Cas9 cloning and targeting strategy was performed as previously described ^41^. sgRNA guide sequence (GTCGTTTGCCTCCATTTCTGT) was chosen to target the TIP5 locus on exon 3 three base pairs upstream of the ATG start codon and was cloned into pSpCas9(BB)-2A-GFP (PX458, Addgene). This plasmid was co-transfected with the HDR repair template plasmid containing the FLAG/HA inclusion flanked by 1kb homology arms into wild type ESCs at a molar ratio of 1:3. After two days, positively transfected cells were selected for GFP expression and were then further cultured for additional three days. Subsequently, ESCs were seeded for single cell clone isolation. Derived clones were genotyped by PCR using two different primer pairs. One PCR aimed to identify site-specific FLAG-HA inclusion, while the other PCR allowed distinguishing between inclusions in one or both alleles (as illustrated in Supplementary Fig.S4). The integrity of the FLAG-HA inclusion was verified by cloning the exon 3 sequences into CloneJET PCR cloning kit (Thermo Scientific) and by Sanger sequencing (Microsynth). FLAG/HA-TIP5 ESC and ESC-wt behave similarly in terms of TIP5 expression levels, lack of TIP5 association with rRNA genes in ESC, proliferation, and expression of pluripotency markers.

### Chromatin immunoprecipitation and mass spectrometric analyses

Approximately 10^8^ ESCs were collected by scraping followed by washing with PBS. Nuclei were isolated by re-suspending the cells in two consecutive rounds in hypotonic buffer (10mM Hepes pH 7.6, 1.5mM MgCl_2_, 10mM KCl, 2mM Na_3_VO_4_ freshly supplemented with cOmplete^™^ Protease Inhibitor Cocktail (Roche)). The suspension was passed through a douncer homogenizer with a loose pestle 10-20 times and the purity of nuclei was checked under a microscope. The chromatin was then isolated and crosslinked by resuspending the nuclei in the chromatin fractionation/crosslinking buffer (10mM Hepes pH 7.6, 3mM MgCl_2_, 150mM NaCl, 0.5% Triton X-100, 0.5mM dithiobis[succinimidylpropionate] (DSP, Thermo Scientific), 2mM Na_3_VO_4_ supplemented with cOmplete^™^ Protease Inhibitor Cocktail (Roche)) and rotation at room temperature for 30min. The crosslinking was stopped by the addition of 25mM Tris-HCl pH 7.5. The chromatin was then isolated by centrifugation and washed twice in MNase digestion buffer (0.3M Sucrose, 50mM Tris-HCl pH 7.5, 30mM KCl, 7.5mM NaCl, 4mM MgCl_2_, 1mM CaCl_2_, 0.125% NP-40, 0.25% NaDeoxycholate, 2mM Na_3_VO_4_ supplemented with cOmplete^™^ Protease Inhibitor Cocktail (Roche)). Digestion of chromatin into mononucleosomes was assured by digestion with 100U MNase (Roche) in MNase digestion buffer at 37°C for 1h. SDS was then added to a final concentration of 1% followed by a 3x 30sec sonication steps with a bioruptor sonicator (Diagenode). Insoluble precipitates were removed by centrifugation and soluble crosslinked chromatin extracts were diluted 10x in IP buffer (0.3M Sucrose, 50mM Tris-HCl pH 7.5, 30mM KCl, 300mM NaCl, 4mM MgCl_2_, 1mM CaCl_2_, 0.125% NP-40, 0.25% NaDeoxycholate, 2mM Na_3_VO_4_ supplemented with cOmplete^™^ Protease Inhibitor Cocktail (Roche)) and 30µl ANTI-FLAG M2 Affinity Gel (Sigma) were added to the extracts. Binding of FLAG/HA-TIP5 was performed by incubation over night at 4°C while rotating. The beads were subsequently washed five times in wash buffer (20mM Tris pH 7.5, 20% glycerol, 100mM KCl, 300mM NaCl, 1.5mM MgCl2, 0.2mM EDTA, 0.125% NP40, 0.25% NaDeoxycholate, 2mM Na_3_VO_4_ supplemented with cOmplete^™^ Protease Inhibitor Cocktail (Roche)). Purified complexes were then eluted with 2mM FLAG peptide (Sigma) in TBS buffer (50mM Tris-HCl pH 8.0, 150mM NaCl). Eluted proteins were precipitated with the addition of 0.25x volume of 100% trichloric acid (Sigma). Protein pellets were washed five times with cold Acetone (Merck) and submitted for subsequent mass spectrometric analyses by the Functional Genomics Center Zurich (FGCZ). The dry pellets were dissolved in 45 µl buffer (10 mM Tris + 2 mM CaCl_2_, pH 8.2) and 5 µl of trypsin (100 ng/µl in 10 mM HCl) for digestion, which was carried out in a microwave instrument (Discover System, CEM) for 30 min at 5 W and 60 °C. Samples were dried in a SpeedVac (Savant). For LC-MS/MS analysis the samples were dissolved in 0.1% formic acid (Romil)) and an aliquot ranging from 5 to 25% was analyzed on a nanoAcquity UPLC (Waters Inc.) connected to a Q Exactive mass spectrometer (Thermo Scientific) equipped with a Digital PicoView source (New Objective). Peptides were trapped on a Symmetry C18 trap column (5 µm, 180 µm × 20 mm, Waters Inc.) and separated on a BEH300 C18 column (1.7 µm, 75 µm × 150 m, Waters Inc.) at a flow rate of 250 nl/min using a gradient from 1% solvent B (0.1% formic acid in acetonitrile, Romil)/99% solvent A (0.1% formic acid in water, Romil) to 40% solvent B/60% solvent A within 90 min. Mass spectrometer settings were: Data dependent analysis. Precursor scan range 350 – 1500 m/z, resolution 70’000, maximum injection time 100 ms, threshold 3e6. Fragment ion scan range 200 – 2000 m/z, Resolution 35’000, maximum injection time 120 ms, threshold 1e5. Proteins were identified using the Mascot search engine (Matrix Science, version 2.4.1). Mascot was set up to search the SwissProt database assuming the digestion enzyme trypsin. Mascot was searched with a fragment ion mass tolerance of 0.030 Da and a parent ion tolerance of 10.0 PPM. Oxidation of methionine was specified in Mascot as a variable modification. Scaffold (Proteome Software Inc.) was used to validate MS/MS based peptide and protein identifications. Peptide identifications were accepted if they achieved a false discovery rate (FDR) of less than 0.1% by the Scaffold Local FDR algorithm. Protein identifications were accepted if they achieved an FDR of less than 1.0% and contained at least 2 identified peptides.

### Chromatin immunoprecipitation (ChIP)

ChIP analysis was performed as previously described ^25^. Briefly, 1% formaldehyde was added to cultured cells to cross-link proteins to DNA. For histone ChIPs, isolated nuclei were then lysed and sonicated using a Bioruptor ultrasonic cell disruptor (Diagenode) to shear genomic DNA to an average fragment size of 200 bp. 20 µg of chromatin was diluted to a total volume of 500 µl with ChIP buffer (16.7 mM Tris–HCl, pH 8.1, 167 mM NaCl, 1.2 mM EDTA, 0.01% SDS, 1.1% Triton X-100) and pre-cleared with 10 µl packed Sepharose beads for 2 h at 4°C. Pre-cleared chromatin was incubated overnight with the indicated antibodies. The next day, Dynabeads protein-A (or -G, Millipore) were added and incubated for 4 h at 4°C. After washing, bound chromatin was eluted with the elution buffer (1% SDS, 100 mM NaHCO_3_). Upon proteinase K digestion (50°C for 3 h) and reversion of cross-linking (65°C, overnight), DNA was purified with phenol/chloroform, ethanol precipitated and quantified by qPCR using the primers listed in **Supplementary Table 7**.

For TIP5 ChIPs, we noticed that sonication of formaldehyde-crosslinked chromatin induced degradation of TIP5 (data not shown). Therefore, to increase the efficiency of TIP5 ChIPseq, we fragmented crosslinked chromatin into mono-nucleosomes through digestion with MNase. Briefly, isolated and crosslinked nuclei were MNase digested in 400µl MNase digestion buffer (0.3M Sucrose, 50mM Tris pH 7.5, 30mM KCl, 7.5mM NaCl, 4mM MgCl_2_, 1mM CaCl_2_, 0.125% NP-40, 0.25% NaDeoxycholate, 2mM Na_3_VO_4_ supplemented with cOmplete^™^ Protease Inhibitor Cocktail (Roche)) with 100U MNase (Roche) at 37°C for 1h. The digestion was then stopped with 5mM EDTA and the digested chromatin was solubilized in 1% SDS and three pulses of 30sec sonication using a Bioruptor ultrasonic cell disruptor (Diagenode). 200µg of pre-cleared chromatin was immunopurified with incubation of 30µl of Anti-FLAG M2 Affinity Gel (Sigma) or 30µl Anti-HA magnetic beads (Pierce) over night. The samples were subsequently washed, eluted and the DNA was purified as for histone ChIPs. ChIP-qPCR measurements were performed with KAPA SYBR^®^ FAST (Sigma) on a Rotor-Gene Q (Qiagen) always comparing enrichments over input samples. Primer sequences are listed in **Supplementary Table 7**.

For ChIPseq analyses, the quantity and quality of the isolated DNA was determined with a Qubit® (1.0) Fluorometer (Life Technologies, California, USA) and a Bioanalyzer 2100 (Agilent, Waldbronn, Germany). The Nugen Ovation Ultra Low Library Systems (Nugen, Inc, California, USA) was used in the following steps. Briefly, ChIP samples (1 ng) was end-repaired and polyadenylated before the ligation of Illumina compatible adapters. The adapters contain the index for multiplexing. The quality and quantity of the enriched libraries were validated using Qubit® (1.0) Fluorometer and the Bioanalyzer 2100 (Agilent, Waldbronn, Germany). The libraries were normalized to 10nM in Tris-Cl 10 mM, pH8.5 with 0.1% Tween 20. The TruSeq SR Cluster Kit v4-cBot-HS (Illumina, Inc, California, USA) was used for cluster generation using 8 pM of pooled normalized libraries on the cBOT. Sequencing was performed on the Illumina HiSeq 2500 single end 126 bp using the TruSeq SBS Kit v4-HS (Illumina, Inc, California, USA).

### ChIPseq data analysis

Own and published ChIPseq reads were aligned to the mouse mm10 reference genome using Bowtie2 (version 2.2.5; ^42^). Read counts were computed and normalized using “bamCoverage” from deepTools (version 2.0.1; ^43^) using a bin size of 50bp. deepTools was also further used to generate all heat maps, profiles and pearson correlation plots. TIP5 bound regions were defined using SICER (version 1.1; ^44^) by comparing the FLAG ChIPs of tagged vs non-tagged TIP5 ESCs in 2i and serum using the following arguments: W=1000 G=3000 FDR=0.00001. These analyses revealed 8824 and 7221 TIP5-bound regions in 2i and serum ESCs, respectively. Highly similar results were obtained by defining TIP5 bound regions comparing the FLAG ChIPs to the input samples, excluding strong biases of the FLAG antibody.

Quantitative ChIPseq experiments were performed by spike-in ESC chromatin with 50ng of *Drosophila* chromatin in order to compare the differences in H3K27me3 levels. ChIP experiments were performed as described above. After sequencing, reads uniquely aligned were mapped to *Drosophila* genome (dm6) and an internal normalization factor was generated for each sample according to Active Motif spike-in normalization strategy. The number of reads for ESC ChIPs were normalized by down-sampling the reads based on the spike-in normalization factor using “bamCoverage” from deepTools. CTCF ChIPseq data sets in 2i and serum ESCs were taken from ^45^. CTCF peaks were defined using MACS2 (version 2.1.0; ^46^) comparing the CTCF ChIPseq to its respective input sample with a qValue cutoff of 0.0001. Using these parameters 56218 and 47245 CTCF peaks were defined in 2i and serum ESCs, respectively. The ratio of mean H3K27me3 read counts 2.5kb upstream and downstream of each CTCF peak were calculated and served as “insulation scores”. H3K27me3-insulated CTCF peaks were defined with an insulation score of >2 or <0.5. Regions with an overall average of less than 0.3 normalized read counts were excluded. This filtering resulted in 7547 CTCF peaks (13.4% of all CTCF peaks) and 8448 peaks (17.9% of all CTCF peaks) that were H3K27me3-insulated in 2i and serum ESCs, respectively. The accuracy of these calculations was confirmed by plotting published H3K27me3 ChIPseq data sets from wild type 2i and serum ESCs ^16^ over these H3K27me3-insulating CTCF peaks revealing highly similar results (data not shown). Pearson correlation plots were generated using deepTools (version 2.0.1). After removal of blacklist regions, chromosome 19 was partitioned into 1kb windows and correlation plots were computed with indicated data sets. H3K36me3 (GSM590119), H3K9me3 (GSM850407) ^16^ and H3K4me1 (GSM1856424) ^17^ were taken from published ChIPseq data sets of ESCs in 2i. For data analysis over transcribed regions, genomic coordinates from all refseq transcripts were retrieved from Ensembl biomart. After removal of blacklist regions, normalized ChIPseq data sets were plotted either over +/− 5kb from the TSS or over the entire transcribed region by scaling the gene length to 20kb (+5kb from TES and −5kb from the TSS). For distinguishing H3K27me3-high and H3K27me3-low promoters, mean read counts +/− 5kb from each TSS were computed. The first quartile was termed as H3K27me3-high, while quartiles 2-4 were defined as H3K27me3-low. TIP5-bound genes were defined by an overlap of a TIP5-bound region with the transcribed regions of the respective gene using bedtools (version 2.24.0; ^47^). Conversion of mm9 and mm10 data sets was performed using Crossmap (version 0.2.4; ^48^). Integrative Genome Viewer (IGV, version 2.3.92) ^49^ was used to visualize and extract representative ChIPseq tracks.

### RNAseq and data analysis

Total RNA from three independent siRNA-mediated TIP5 knockdown experiments was purified with TRIzol reagent (Life Technologies) as stated above. In order to remove DNA contaminants, the samples were treated with 1U DNaseI (Thermo Scientific) for 1h at 37°C and the RNA samples were re-purified using TRIzol. The quality of the isolated RNA was determined with a Qubit® (1.0) Fluorometer (Life Technologies, California, USA) and a Bioanalyzer 2100 (Agilent, Waldbronn, Germany). Only those samples with a 260 nm/280 nm ratio between 1.8–2.1 and a 28S/18S ratio within 1.5–2 were further processed. The TruSeq RNA Sample Prep Kit v2 (Illumina, Inc, California, USA) was used in the succeeding steps. Briefly, total RNA samples (100-1000 ng) were poly A enriched and then reverse-transcribed into double-stranded cDNA. The cDNA samples was fragmented, end-repaired and polyadenylated before ligation of TruSeq adapters containing the index for multiplexing Fragments containing TruSeq adapters on both ends were selectively enriched with PCR. The quality and quantity of the enriched libraries were validated using Qubit® (1.0) Fluorometer and the Caliper GX LabChip® GX (Caliper Life Sciences, Inc., USA). The product is a smear with an average fragment size of approximately 260 bp. The libraries were normalized to 10nM in Tris-Cl 10 mM, pH8.5 with 0.1% Tween 20. The TruSeq SR Cluster Kit HS4000 (Illumina, Inc, California, USA) was used for cluster generation using 10 pM of pooled normalized libraries on the cBOT. Sequencing was performed on the Illumina HiSeq 4000 single end 100 bp using the TruSeq SBS Kit HS4000 (Illumina, Inc, California, USA). Reads were aligned to the reference genome (ensembl version 82) with Subread (i.e. subjunc, version 1.4.6-p4; ^50^) allowing up to 16 alignments per read (options: –trim5 10 –trim3 15 -n 20 -m 5 -B 16 -H –allJunctions). Count tables were generated with Rcount ^51^ with an allocation distance of 100 bp for calculating the weights of the reads with multiple alignments, considering the strand information, and a minimal number of 5 hits. Variation in gene expression was analyzed with a general linear model in R with the package edgeR (version 3.12.0; ^52^) according to a crossed factorial design with two explanatory factors (i) siRNA against Tip5 and a mock sequence and (ii) ESCs grown in 2i or serum. Genes differentially expressed between specific conditions were identified with linear contrasts using trended dispersion estimates and Benjamini-Hochberg multiple testing corrections. Genes with a *P-*value below 0.05 and a minimal fold change of 1.5 were considered to be differentially expressed. These thresholds have previously been used characterizing chromatin remodeler functions ^53^. Gene ontology analysis was performed with David Bionformatics Resource 6.8 ^54^.

### ATACseq analysis

ATAC-seq experiments were performed in biological triplicates of mES cells grown in 2i or serum conditions. Cell pellets from 50,000 cells for each condition were freshly harvested 72 h after transfection with siRNAs against TIP5 or a control sequence. The tagmentation reaction was performed as previously described ^55^ with minor adjustments of the protocol, using the Nextera DNA Library Prep Kit (Illumina) together with the barcoded primers from the Nextera Index Kit (Illumina). In brief, an additional size selection step was performed after the first 5 cycles of library amplification. For this, the PCR reaction was incubated with 0.6 × volume of Ampure XP beads (Beckman Coulter) for 5 min to allow binding of high molecular weight fragments. Beads containing long DNA fragments were separated on a magnet and the supernatant containing only small DNA fragments below roughly 800 bp were cleaned up using MinElute PCR purification columns (Qiagen). All libraries were amplified for 12 cycles in total, visualized and quantified with a TapeStation2200 (Agilent), and sequenced on a Illumina HiSeq4000 instrument obtaining 125 bp single-end reads.

Raw sequencing reads were filtered for low-quality after adapter removal and aligned on mm9 using Bowtie2 with “-- very sensitive” mapping parameters. PCR duplicates were removed using Picard and only uniquely mapped reads were considered for further analysis. Due to very low transposition efficiency, ESC+serum/siControl replicate 2 was excluded from any further analysis. To call differentially open chromatin sites between the various conditions, the mapped reads of all conditions were merged into one alignment file. Peaks were called on the merged sample using MACS2 with -- nomodel --shift -100 --extsize 200 --keep-dup all, to obtain a consensus peak set with equal contribution of each sample independent of the respective biological treatment ^56^. The combined peak set was used to build a count matrix using the aligned reads of the individual samples. Differential peaks between sample groups were identified using edgeR ^57^. For this, the count matrix was filtered for peaks with low coverage across all samples based on a average CPM value < −1, resulting in a final set of 128,253 peaks with p-values below 0.0034. The remaining counts were normalized using the total library size as well as edgeR’s TMM derived normalization factors. Between two sample groups, only regions with a log fold change above > 0.5 and a FDR value < 0.1 and group_CPM > −0.2 were considered as differentially accessible for further analysis. Identified ATAC peaks were overlapped with known genomic elements using bedtools (version 2.24.0; ^47^ with a minimum of one bp overlap from both elements. Annotation of gained and lost ATAC peaks upon TIP5 knockdown in ESC+2i was performed using ChIPseeker ^58^.

### HiC and data analysis

HiC experiments were performed in triplicates of ESC+2i treated with siRNA-control or siRNA-*TIP5*. We generated roughly 200 million valid pair end reads from ESC+2i treated with siRNA-control (95 million) and siRNA-*Tip5* (105 million).

Five million cells were pelleted and resuspended in PBS-10%FCS. PBS-10%FCS-4% formaldehyde was added to a final concentration of 2% formaldehyde (v/v). Samples were incubated at room temperature for 10 minutes with mixing. Ice-cold glycine solution was added to a final concentration of 0.2M and immediately centrifuged for 5 minutes at 300xg at 4°C. Cells were washed in 1 ml of ice-cold PBS. Pellet was flash-frozen in liquid nitrogen. Pellet was taken up and washed in 1 ml ice cold lysis buffer1, resupended again in 1 ml of ice cold lysis buffer and incubated for 30 minutes at 4°C. Sample was pelleted and washed in 0.5 ml 1.2x DpnII buffer, resuspended in 0.5 ml of DpnII buffer again and moved to a thermomixer at 37°C and 300 rpm. SDS was then carefully added to a concentration of 0.3%, slowly suspended with a pipet and incubated for an hour. Triton X-100 was added to a concentration of 2.6% and sample was incubated for another hour. To digest the sample, 200 units of DpnII enzyme were added for a 4 hour incubation in a thermomixer at 37°C and 900 rpm; another 200 units of DpnII were added for overnight incubation. From here on, we adopted the protocol described in ^2^ with some adjustments Cells were incubated for 20 minutes at 65°C to heat inactivate DpnII, pelleted and 300 µl of fill-in mastermix was added (218 µl of MilliQ, 30 µl of 10x NEB buffer 2, 15 µl of 10mM dCTP, 15 µl of 10mM dGTP, 15 µl of 10mM dTTP, 37.5 µl of 0.4mM biotin-14-dATP (Life Technologies, 19524-016), 10 µl of 5U/µl DNA Polymerase I, Large (Klenow) Fragment (NEB, M0210)). Sample was mixed by pipetting and incubated for 60 minutes at 37°C shaking at 300 rpm and placed at 4°C afterwards. 900 µl of ligation mix was added (120 µl 10x ligase buffer, 50 units of T4 ligase (Roche) and 770 µl MilliQ) and mixed by inverting and incubated overnight at 16°C. Sample was pelleted for 5 minutes at 1000 × g and taken up in 500 µl 10 mM Tris. Protein was degraded by adding 50 µl of 20mg/ml proteinase K (NEB, P8102), 50 µl of 10% SDS and incubated at 55°C for 30 minutes. 57 µl of 5M of sodium chloride was then added and the sample was incubated at 68°C overnight or for at least 1.5 hours. Samples were cooled to room temperature and DNA was purified using NucleoMag P-Beads and taken up in 5mM Tris pH7.5. Samples were sheared to a size of 300-500 bp using a Covaris S2 focused-ultrasonicator. From here on, the protocol described in ^2^ was adopted.

FastQ files were mapped to the mouse genome (mm10) using bwa-mem ^59^ and filtered and deduplicated using HiCUP v0.5.10 ^60^. Chromosomal interaction matrices were generated using Juicer ^61^ at 500 Kb resolution and normalized by Knight and Ruiz’s matrix balancing algorithm. Biological replicates were first processed independently and inspected for clustering between TIP5 depletion and control conditions by PCA. Next, replicates per condition were pooled to create merged contact maps that were used in the downstream analyses.

Eigenvalues of samples calculated using Juicer Eigenvector were used to annotated A and B genome compartments and TADs were calculated using Juicer Arrowhead ^1^. To visualize the impact of Tip5 knockdown in chromosomal architecture, we plotted the median contact frequency from each genomic region at increasing genomic distance. ENCODE Data Analysis Consortium Blacklisted Regions ^62^ were excluded from the analysis.

Enrichment or depletion of genomic features was tested, relative to expectation, at subregions within genome compartments, defined by dividing the compartments into 5 equal size segments where the two outer bins represent compartment boundaries and the central bin as compartment center, using the Genome Association Tester (GAT) ^63^. Specifically, sub-compartment enrichment was compared to a null distribution obtained by randomly sampling 10,000 times (with replacement) segments of the same length and matching GC content as the tested feature within all annotated compartments. The genome was also divided into segments of 10 Kb and assigned to eight isochore bins to control for potential confounding variables that correlate with GC content, such as gene density, in the enrichment analysis.

Genome-wide Pearson correlation of own and published ChIPseq data sets with high resolution HiC eigenvector values ^8^ were calculated genome-wide at 5kb resolution using deeptools (version 2.0.1; ^43^). Published ChIPseq data sets were taken for CTCF GSM2259907 ^64^, H3K9me2 GSM2051614 ^65^, H3K9me3 GSM850406, H3K36me3 GSM590119 ^16^, H3K27ac GSM1856425, H3K4me1 GSM1856423, Ring1b GSM1856437 ^17^, TOP2A GSM1110842 ^66^ and Smc3 GSM560343 ^67^.

## Public Datasets used in this study

Public datasets used in this study are listed in **Supplementary Table 7**.

## Accession numbers

All raw data generated in this study using high throughput sequencing are accessible through NCBI’s GEO (accession number GSE112222).

## Acknowledgements

This work was supported by the Swiss National Science Foundation (310003A-152854 and 31003A_173056 to R.S.; PP00P3_150667 and NCCR RNA & Disease to ACM; 157488 and 180345 to T.B.), ERC grant (ERC-AdG-787074-NucleolusChromatin to RS), Forschungskredit of the University of Zurich (to D.D and E.V.), UBS Promedica Stiftung, Julius Müller Stiftung, Olga Mayenfisch Stifung and Stiftung für wissenschaftliche Forschung an der Universität Zürich (to R.S.). We thank Peter Hunziker, Catherine Aquino and the Functional Genomic Center Zurich for the assistance in sequencing and proteomic analysis. We also thank Dominik Bär for technical assistance. We thank C. Ciaudo for having provided ESC lines.

## Supplementary Figures

**Supplementary Figure 1**

**TIP5 knockdown affects proliferation of ESC+2i**

**a.** Data represent relative cell numbers and were normalized to ESC transfected with siRNA-Control. TIP5 knockdown was achieved with a different siRNA-*Tip5* sequence (siRNA-*Tip5#2*). Average values of three independent experiments. Error bars represent s.d.

**b.** Data represent relative cell numbers using another ESC line (ESC#2) cultured in 2i medium and were normalized to ESC transfected with siRNA-Control. Average values of two independent experiments.

**c.** Flow cytometry (FACS) profile of cell cycle progression of ESC+2i and ESC+serum upon treatment with siRNA-Control and siRNA-*Tip5* and its quantification (**d**) from two independent experiments.

**e.** Western blot showing absence of apoptotic markers in ESC+2i upon treatment with siRNA-*Tip5*. ESC+2i treated with staurosporine represent the positive control for apoptotic markers signals PARP1 and Caspase 3 cleavage.

**f.** TIP5-KO in ESC+serum does not affect differentiation. Representative images of ESC and cells after 3 days of differentiation.

**g.** Data represent relative cell numbers of ESC and Tip5-KO ESC+serum after transition from serum to 2i conditions and were normalized to ESC cultured in serum.

**Supplementary Figure 2**

**a.** Venn diagrams showing number of differently expressed genes upon TIP5 knockdown detected in ESC+2i compared to ESC+serum.

**b.** qRT-PCR. Expression analysis of TIP5 regulated genes in ESC+2i upon TIP5 knockdown using a different siRNA-*Tip5* (siRNA-*Tip5#2*). Upregulated genes in ESC+2i upon TIP5-KD are labeled in blue, downregulated genes are in red. mRNA levels were normalized to *Rps12* mRNA and to ESC+2i transfected with siRNA-Control. Average values of three independent experiments. Error bars represent s.d. Statistical significance (*P*-values) for the experiments was calculated using the paired two-tailed t-test (ns: non significant, *<0.05, **< 0.01, ***<0.001, ****<0.0001).

**c.** qRT-PCR. Expression analysis of TIP5 regulated genes in another ESC line (ESC#2) cultured in 2i medium upon TIP5 knockdown. mRNA levels were normalized to *Rps12* mRNA and to ESC+2i transfected with siRNA-Control. Experiment performed as one replicate.

**d.** TIP5 knockdown does not affect proliferation of TKO-ESCs cultured in serum/LIF. Data represent relative cell numbers after 3 days of siRNA treatment and were normalized to ESC transfected with siRNA-Control. Average values of three independent experiments. Error bars represent s.d. Statistical significance (*P*-values) for the experiments was calculated using the paired two-tailed t-test (ns: non significant).

**e.** Knockdown of TIP5 in TKO-ESCs does not affect transcription of genes regulated by TIP5 in ESC+2i. mRNA levels were normalized to *Rps12* mRNA and to ESCs transfected with siRNA-Control. Average values of three independent experiments. Error bars represent s.d. Statistical significance (*P*-values) for the experiments was calculated using the paired two-tailed t-test (*<0.05, ****<0.0001). Data without P values are statistically non significant.

**f.** TIP5 knockdown does not affect proliferation of Eed^−/−^ and Ring1b^−/−^ ESCs cultured in serum. Data represent relative cell numbers after 3 days of siRNA treatment and were normalized to ESC transfected with siRNA-Control. Average values of three independent experiments. Error bars represent s.d. Statistical significance (*P*-values) for the experiments was calculated using the paired two-tailed t-test (***<0.0001). Data without P values are statistically non significant.

**g.** TIP5 knockdown does affect expression of genes regulated by TIP5 in Eed^−/−^ and Ring1b^−/−^ ESCs cultured in serum. RT-qPCR experiments in Eed^−/−^ (left panel) and Ring1b^−/−^ (right panel) -ESCs cultured in serum treated with siRNA-control or siRNA-*Tip5.* mRNA levels were normalized to *Rps12* mRNA and ESC+2i transfected with siRNA-Control. Average values of three independent experiments. Error bars represent s.d. Statistical significance (*P*-values) for the experiments was calculated using the unpaired two-tailed t-test (ns: non significant, ***<0.001).

**h.** Boxplot showing that the mean of normalized H3K27me3 reads at TIP5-regulated and random gene promoters (+/− 1kb) in ESC+serum. Statistical significance (*P*-values) for the experiments was calculated using the unpaired two-tailed t-test (*** < 0.001).

**i.** Boxplot showing log2 fold-changes of normalized H3K27me3 read counts at TIP5-regulated and random gene promoters (+/− 1kb) in ESC+serum relative to ESC+2i.

**Supplementary Figure 3**

**a.** Scheme representing the strategy to insert FLAG-HA sequences at the 5’ of the endogenous TIP5 sequence.

**b, c.** PCR genotyping of four clones for insertion of the F/H-TIP5 sequence on the *TIP5* locus. Clones #1 and #3 were homozygous for *F/H-TIP5* alleles. Sanger sequencing confirmed the integrity of F/H-TIP5.

**d.** Representative image showing that TIP5 associates with internal chromatin domains (ICD) in ESC+2i and ESC+serum. The data refer to TIP5 ChIPseq performed with FLAG-antibodies with *F/H-Tip5* ESC+2i and ESC+serum. Data for LADs and ICDs refer to ESC+serum and are from ^28^.

**e.** Venn diagrams showing TIP5 bound regions in ESC+2i and ESC+serum.

**f.** TIP5-bound enhancers are not linked to TIP5-regulated genes in linear distance. The distances of all TIP5-bound enhancers and enhancers bound by TIP5 only in ESC+2i to the nearest gene promoter (+/− 5kb of TSS) were calculated. Log10 distances in bp to the nearest upregulated, downregulated and not regulated gene promoter are plotted.

**g.** TIP5-regulated genes are not linked to TIP5-bound enhancers in linear distance. The distances of TIP5-regulated gene promoters (+/− 5kb of TSS) to all TIP5-bound enhancers, enhancers bound by TIP5 only in ESC+2i and enhancers not bound by TIP5 were calculated. Log10 distances in bp are plotted.

**Supplementary Figure 4**

**A.** Boxplots indicating observed/expected contact values between TIP5-bound regions and loci not bound by TIP5 at near-*cis* contacts in all genome or in A compartments of ESC+serum, NPC and CN.

**b.** Histogram of the coverage of A and B compartments annotations from HiC of this study in ESC+2i compared to those annotated with HiC of ESC+serum generated by ^8^.

**c.** Both ESC+2i and ESC+serum show strong long-distance contacts within A compartments. Plot showing the mean of observed/expected contact values relative to genomic distance between A-to-A, A-to-B and B-to-B compartment contacts. Values from ESC+serum were from ^8^.

**d.** Long distance contacts and compartmentalization is similar in ESC+2i and ESC+serum. Hi-C contact matrices for a zoomed in region on chromosome 6 at 100-kb resolution showing the presence of strong long-distance contacts corresponding to regions in A compartment bound by TIP5 in ESC+2i. This image should be compared with the image of ESC+serum shown in Figure 3b.

**e.** TIP5 binding marks the A subcompartment. Boxplot indicating the eigenvector values from high resolution HiC data in ESC+serum ^8^ and ESC+2i that were calculated from the top of TIP5-bound regions and compared to the same number of random regions within the A compartment that were not bound by TIP5. Statistical significance (*P*-values) for the experiments was calculated using the unpaired two-tailed t-test (*** < 0.001).

**Supplementary Figure 5**

**a.** Venn diagrams showing ATACseq peaks in ESC+2i and ESC+serum.

**b.** Numbers of ATAC peaks called separately in ESC+2i and ESC+serum and their association with TIP5.

**c.** Representative images showing increased chromatin accessibility in ESC+2i upon TIP5 depletion.

**d.** TIP5 depletion increases chromatin accessibility only in ESC+2i. Numbers of significant changed ATAC peaks in ESC+2i and ESC+serum upon TIP5 knockdown including their association with TIP5.

**Supplementary Figure 6**

**a.** PCA analysis of HiC replicates of ESC+2i treated with siRNA-control and siRNA-*Tip5*.

**b.** Relative contact frequencies over genomic distances in ESCs of replicates treated with siRNA-control and siRNA-TIP5.

**c.** Relative contact frequencies over genomic distances of different genomic regions in ESCs treated with siRNA-control and siRNA-TIP5.

**Supplementary Figure 7**

**a.** Western blot showing total levels of H3K27me3, H3K27ac and H3K4me3 in ESC+2i transfected with siRNA-control and siRNA-*Tip5*. Histone H3 is shown as a protein loading control.

**b.** Quantitative H3K27me3 ChIP **using** *Drosophila* spike-in. Scatterplot showing the changes in H3K27me3 upon TIP5 depletion in ESC+2i and ESC+serum. Mean of normalized H3K27me3 read counts in ESC+2i and ESC+serum treated with siRNA-control or siRNA-*Tip5* over all TSSs (+/− 1kb) were calculated. Log2-fold changes of (siRNA-*Tip5/*siRNA-control) are plotted relative to the mean of normalized H3K27me3 occupancies.

**c.** H3K27me3 ChIP in ESC+2i upon TIP5 knockdown using a different siRNA-*Tip5* (siRNA-*Tip5#2*). Values show H3K27me3 occupancy at promoters of genes regulated by TIP5 and at four sequences (#1-#4) with elevated H3K27me3 at CTCF boundaries (H3K27me3^hi^/CTCF). Data were measured by qPCR and normalized to input and *Tshz1* value in control cells. Average values of three independent experiments. Error bars represent s.d.

**d.** TIP5 knockdown affects proliferation of Eed^−/−^ and Ring1b^−/−^ ESCs cultured in 2i. Data represent relative cell numbers after 3 days of siRNA treatment and were normalized to ESC transfected with siRNA-Control. Average values of three independent experiments. Error bars represent s.d. Statistical significance (*P*-values) for the experiments was calculated using the unpaired two-tailed t-test (**<0.01).

**e.** TIP5 depletion in ESC+2i also affects gene expression in the absence of PRC-mediated repression. RT-qPCR experiments in Eed^−/−^ and Ring1b^−/−^-ESCs cultured in 2i treated with siRNA-control or siRNA-*Tip5.* Upregulated genes in ESC+2i upon TIP5-KD are labeled in blue, downregulated genes are in red. mRNA levels were normalized to *Rps12* mRNA and ESC+2i transfected with siRNA-Control. Average values of three independent experiments. Error bars represent s.d. Statistical significance (*P*-values) for the experiments was calculated using the paired two-tailed t-test (*<0.05, **< 0.01, ***<0.001). Data without P values are statistically non significant.

**f.** Hi-C contact matrices for a zoomed in region on chromosome 11 containing *HoxB* cluster (5 kb resolution). Blue arrows indicated gain or increase of contacts in ESC+siRNA-*Tip5* compared to control cells (siRNA-control). Orange arrows indicate loss or decrease in contacts.

**Supplementary Figure 8**

**A.** SMC3 and TIP5 regulate gene expression in ESC+2i via a shared pathway. qRT-PCR of genes regulated by TIP5 in ESC+2i (upper panel) and in ESC+serum (rbottom panel) upon SMC3 or TIP5 knockdown by siRNA. Upregulated genes in ESC+2i upon TIP5-KD are labeled in blue, downregulated genes are in red. mRNA levels were normalized to *Rps12* mRNA and to ESC transfected with siRNA-Control. Average values of three independent experiments. Error bars represent s.d. Statistical significance (*P*-values) for the experiments was calculated using the paired two-tailed t-test (*<0.05, **< 0.01, ***<0.001). Data without P values are statistically non significant.

**b.** Depletion of Smc3 does not affect H3K27me3 occupancy in ESC+2i. H3K27me3 ChIP values represent the average values of three independent experiments. Error bars represent s.d. Statistical significance (*P*-values) for the experiments was calculated using the paired two-tailed t-test (ny: non significant, *<0.05, **< 0.01, ***<0.001).

**c.** TIP5 binding in ESC+2i depends on TOP2A. Anti-FLAG ChIP in F/H-TIP5 ESCs treated with siRNA-control or siRNA-*Top2a*. Values from two independent experiments were normalized to their respective input and to ATF7IP in control cells.

**d.** TIP5 binding in ESCs depends on TOP2A activity. Anti-FLAG ChIP in F/H-TIP5-ESCs cultured in 2i (three experiments) or serum (two experiments) treated for 24 hours with DMSO or ICF-193. Values were normalized to to their respective input and to ATF7IP in control cells. In the experiment with ESC+2i average values are from three independent experiments and error bars represent s.d. Statistical significance (*P*-values) for the experiments was calculated using the paired two-tailed t-test (ns: non significant, *<0.05, **< 0.01).

**e.** TIP5 binding in ESC+2i depends on cohesin. Anti-FLAG ChIP in F/H-TIP5-ESCs cultured in 2i or serum treated with siRNA-control or siRNA-*Smc3*. Values were normalized to input and to ATF7IP in control cells. Data are from two independent experiments.

**f.** The association of cohesin with chromatin does not depend on TIP5. Anti-SMC1a ChIP in ESC+2i treated with siRNA-*Tip5.* Average values of three independent experiments. Error bars represent s.d. Statistical significance (*P*-values) for the experiments was calculated using the paired two-tailed t-test (ns: non significant.)

## Supplementary Tables

**Supplementary Table 1**

RNAseq analysis of ESCs treated with siRNA-Control and siRNA-*Tip5*.

**Supplementary Table 2**

GO analysis of genes regulated by TIP5 in ESCs.

**Supplementary Table 3**

Enrichment of DHS and TIP5-regulated genes in A and B compartments in ESCs.

**Supplementary Table 4**

QC process of the HiC

**Supplementary Table 5**

HiC contact values (10 kb resolution) at HoxA cluster in ESC+2i treated with siRNA-Control and siTIP5-siRNA. Values referred to genomic contacts highlighted in **Figure 7i**.

**Supplementary Table 6**

List of TIP5 interacting proteins on chromatin of ESCs.

**Supplementary Table 7**

List of materials, software and datasets used in these study.

## Reference

1. Lieberman-Aiden, E., van Berkum, N.L., Williams, L., Imakaev, M., Ragoczy, T., Telling, A., Amit, I., Lajoie, B.R., Sabo, P.J., Dorschner, M.O., Sandstrom, R., Bernstein, B., Bender, M.A., Groudine, M., Gnirke, A., Stamatoyannopoulos, J., Mirny, L.A., Lander, E.S. & Dekker, J. Comprehensive mapping of long-range interactions reveals folding principles of the human genome. Science 326, 289–293 (2009).

2. Rao, S.S., Huntley, M.H., Durand, N.C., Stamenova, E.K., Bochkov, I.D., Robinson, J.T., Sanborn, A.L., Machol, I., Omer, A.D., Lander, E.S. & Aiden, E.L. A 3D map of the human genome at kilobase resolution reveals principles of chromatin looping. Cell 159, 1665–1680 (2014).

3. Nora, E.P., Goloborodko, A., Valton, A.L., Gibcus, J.H., Uebersohn, A., Abdennur, N., Dekker, J., Mirny, L.A. & Bruneau, B.G. Targeted Degradation of CTCF Decouples Local Insulation of Chromosome Domains from Genomic Compartmentalization. Cell 169, 930–944 e922 (2017).

4. Schwarzer, W., Abdennur, N., Goloborodko, A., Pekowska, A., Fudenberg, G., Loe-Mie, Y., Fonseca, N.A., Huber, W., C, H.H., Mirny, L. & Spitz, F. Two independent modes of chromatin organization revealed by cohesin removal. Nature 551, 51–56 (2017).

5. Haarhuis, J.H.I., van der Weide, R.H., Blomen, V.A., Yanez-Cuna, J.O., Amendola, M., van Ruiten, M.S., Krijger, P.H.L., Teunissen, H., Medema, R.H., van Steensel, B., Brummelkamp, T.R., de Wit, E. & Rowland, B.D. The Cohesin Release Factor WAPL Restricts Chromatin Loop Extension. Cell 169, 693–707 e614 (2017).

6. Rao, S.S.P., Huang, S.C., Glenn St Hilaire, B., Engreitz, J.M., Perez, E.M., Kieffer-Kwon, K.R., Sanborn, A.L., Johnstone, S.E., Bascom, G.D., Bochkov, I.D., Huang, X., Shamim, M.S., Shin, J., Turner, D., Ye, Z., Omer, A.D., Robinson, J.T., Schlick, T., Bernstein, B.E., Casellas, R., Lander, E.S. & Aiden, E.L. Cohesin Loss Eliminates All Loop Domains. Cell 171, 305–320 e324 (2017).

7. Dixon, J.R., Jung, I., Selvaraj, S., Shen, Y., Antosiewicz-Bourget, J.E., Lee, A.Y., Ye, Z., Kim, A., Rajagopal, N., Xie, W., Diao, Y., Liang, J., Zhao, H., Lobanenkov, V.V., Ecker, J.R., Thomson, J.A. & Ren, B. Chromatin architecture reorganization during stem cell differentiation. Nature 518, 331–336 (2015).

8. Bonev, B., Mendelson Cohen, N., Szabo, Q., Fritsch, L., Papadopoulos, G.L., Lubling, Y., Xu, X., Lv, X., Hugnot, J.P., Tanay, A. & Cavalli, G. Multiscale 3D Genome Rewiring during Mouse Neural Development. Cell 171, 557–572 e524 (2017).

9. Hackett, J.A. & Surani, M.A. Regulatory principles of pluripotency: from the ground state up. Cell Stem Cell 15, 416–430 (2014).

10. Ying, Q.L., Wray, J., Nichols, J., Batlle-Morera, L., Doble, B., Woodgett, J., Cohen, P. & Smith, A. The ground state of embryonic stem cell self-renewal. Nature 453, 519–523 (2008).

11. Boroviak, T., Loos, R., Bertone, P., Smith, A. & Nichols, J. The ability of inner-cell-mass cells to self-renew as embryonic stem cells is acquired following epiblast specification. Nat Cell Biol 16, 516–528 (2014).

12. Ricci, M.A., Manzo, C., Garcia-Parajo, M.F., Lakadamyali, M. & Cosma, M.P. Chromatin fibers are formed by heterogeneous groups of nucleosomes in vivo. Cell 160, 1145–1158 (2015).

13. Habibi, E., Brinkman, A.B., Arand, J., Kroeze, L.I., Kerstens, H.H., Matarese, F., Lepikhov, K., Gut, M., Brun-Heath, I., Hubner, N.C., Benedetti, R., Altucci, L., Jansen, J.H., Walter, J., Gut, I.G., Marks, H. & Stunnenberg, H.G. Whole-genome bisulfite sequencing of two distinct interconvertible DNA methylomes of mouse embryonic stem cells. Cell stem cell 13, 360–369 (2013).

14. Leitch, H.G., McEwen, K.R., Turp, A., Encheva, V., Carroll, T., Grabole, N., Mansfield, W., Nashun, B., Knezovich, J.G., Smith, A., Surani, M.A. & Hajkova, P. Naive pluripotency is associated with global DNA hypomethylation. Nat Struct Mol Biol 20, 311–316 (2013).

15. Ficz, G., Hore, T.A., Santos, F., Lee, H.J., Dean, W., Arand, J., Krueger, F., Oxley, D., Paul, Y.L., Walter, J., Cook, S.J., Andrews, S., Branco, M.R. & Reik, W. FGF signaling inhibition in ESCs drives rapid genome-wide demethylation to the epigenetic ground state of pluripotency. Cell Stem Cell 13, 351–359 (2013).

16. Marks, H., Kalkan, T., Menafra, R., Denissov, S., Jones, K., Hofemeister, H., Nichols, J., Kranz, A., Stewart, A.F., Smith, A. & Stunnenberg, H.G. The transcriptional and epigenomic foundations of ground state pluripotency. Cell 149, 590–604 (2012).

17. Joshi, O., Wang, S.Y., Kuznetsova, T., Atlasi, Y., Peng, T., Fabre, P.J., Habibi, E., Shaik, J., Saeed, S., Handoko, L., Richmond, T., Spivakov, M., Burgess, D. & Stunnenberg, H.G. Dynamic Reorganization of Extremely Long-Range Promoter-Promoter Interactions between Two States of Pluripotency. Cell Stem Cell 17, 748–757 (2015).

18. Schoenfelder, S., Sugar, R., Dimond, A., Javierre, B.M., Armstrong, H., Mifsud, B., Dimitrova, E., Matheson, L., Tavares-Cadete, F., Furlan-Magaril, M., Segonds-Pichon, A., Jurkowski, W., Wingett, S.W., Tabbada, K., Andrews, S., Herman, B., LeProust, E., Osborne, C.S., Koseki, H., Fraser, P., Luscombe, N.M. & Elderkin, S. Polycomb repressive complex PRC1 spatially constrains the mouse embryonic stem cell genome. Nat Genet 47, 1179–1186 (2015).

19. McLaughlin, K., Flyamer, I.M., Thomson, J.P., Mjoseng, H.K., Shukla, R., Williamson, I., Grimes, G.R., Illingworth, R.S., Adams, I.R., Pennings, S., Meehan, R.R. & Bickmore, W.A. DNA Methylation Directs Polycomb-Dependent 3D Genome Re-organization in Naive Pluripotency. Cell Rep 29, 1974–1985 e1976 (2019).

20. Santoro, R., Li, J. & Grummt, I. The nucleolar remodeling complex NoRC mediates heterochromatin formation and silencing of ribosomal gene transcription. Nature genetics 32, 393–396 (2002).

21. Strohner, R., Nemeth, A., Jansa, P., Hofmann-Rohrer, U., Santoro, R., Langst, G. & Grummt, I. NoRC--a novel member of mammalian ISWI-containing chromatin remodeling machines. EMBO J 20, 4892–4900 (2001).

22. Mayer, C., Schmitz, K.M., Li, J., Grummt, I. & Santoro, R. Intergenic transcripts regulate the epigenetic state of rRNA genes. Mol Cell 22, 351–361 (2006).

23. Guetg, C., Lienemann, P., Sirri, V., Grummt, I., Hernandez-Verdun, D., Hottiger, M.O., Fussenegger, M. & Santoro, R. The NoRC complex mediates the heterochromatin formation and stability of silent rRNA genes and centromeric repeats. The EMBO journal 29, 2135–2146 (2010).

24. Savić, N., Bär, D., Leone, S., Frommel, S.C., Weber, F.A., Vollenweider, E., Ferrari, E., Ziegler, U., Kaech, A., Shakhova, O., Cinelli, P. & Santoro, R. in Cell Stem Cell, Vol. 15 720–7342014).

25. Leone, S., Bar, D., Slabber, C.F., Dalcher, D. & Santoro, R. The RNA helicase DHX9 establishes nucleolar heterochromatin, and this activity is required for embryonic stem cell differentiation. EMBO Rep 18, 1248–1262 (2017).

26. Guetg, C., Scheifele, F., Rosenthal, F., Hottiger, M.O. & Santoro, R. Inheritance of Silent rDNA Chromatin Is Mediated by PARP1 via Noncoding RNA. Mol Cell 45, 790–800 (2012).

27. Savic, N., Bar, D., Leone, S., Frommel, S.C., Weber, F.A., Vollenweider, E., Ferrari, E., Ziegler, U., Kaech, A., Shakhova, O., Cinelli, P. & Santoro, R. lncRNA Maturation to Initiate Heterochromatin Formation in the Nucleolus Is Required for Exit from Pluripotency in ESCs. Cell Stem Cell 15, 720–734 (2014).

28. Peric-Hupkes, D., Meuleman, W., Pagie, L., Bruggeman, S.W., Solovei, I., Brugman, W., Graf, S., Flicek, P., Kerkhoven, R.M., van Lohuizen, M., Reinders, M., Wessels, L. & van Steensel, B. Molecular maps of the reorganization of genome-nuclear lamina interactions during differentiation. Mol Cell 38, 603–613 (2010).

29. Dowen, J.M., Fan, Z.P., Hnisz, D., Ren, G., Abraham, B.J., Zhang, L.N., Weintraub, A.S., Schujiers, J., Lee, T.I., Zhao, K. & Young, R.A. Control of cell identity genes occurs in insulated neighborhoods in mammalian chromosomes. Cell 159, 374–387 (2014).

30. Nitiss, J.L. DNA topoisomerase II and its growing repertoire of biological functions. Nat Rev Cancer 9, 327–337 (2009).

31. Tiwari, V.K., Burger, L., Nikoletopoulou, V., Deogracias, R., Thakurela, S., Wirbelauer, C., Kaut, J., Terranova, R., Hoerner, L., Mielke, C., Boege, F., Murr, R., Peters, A.H., Barde, Y.A. & Schubeler, D. Target genes of Topoisomerase IIbeta regulate neuronal survival and are defined by their chromatin state. Proc Natl Acad Sci U S A 109, E934–943 (2012).

32. Akimitsu, N., Adachi, N., Hirai, H., Hossain, M.S., Hamamoto, H., Kobayashi, M., Aratani, Y., Koyama, H. & Sekimizu, K. Enforced cytokinesis without complete nuclear division in embryonic cells depleting the activity of DNA topoisomerase IIalpha. Genes Cells 8, 393–402 (2003).

33. Pommier, Y., Leo, E., Zhang, H. & Marchand, C. DNA topoisomerases and their poisoning by anticancer and antibacterial drugs. Chem Biol 17, 421–433 (2010).

34. Cabianca, D.S., Munoz-Jimenez, C., Kalck, V., Gaidatzis, D., Padeken, J., Seeber, A., Askjaer, P. & Gasser, S.M. Active chromatin marks drive spatial sequestration of heterochromatin in C. elegans nuclei. Nature 569, 734–739 (2019).

35. Miller, E.L., Hargreaves, D.C., Kadoch, C., Chang, C.Y., Calarco, J.P., Hodges, C., Buenrostro, J.D., Cui, K., Greenleaf, W.J., Zhao, K. & Crabtree, G.R. TOP2 synergizes with BAF chromatin remodeling for both resolution and formation of facultative heterochromatin. Nat Struct Mol Biol 24, 344–352 (2017).

36. Nuebler, J., Fudenberg, G., Imakaev, M., Abdennur, N. & Mirny, L.A. Chromatin organization by an interplay of loop extrusion and compartmental segregation. Proc Natl Acad Sci U S A 115, E6697–E6706 (2018).

37. Uuskula-Reimand, L., Hou, H., Samavarchi-Tehrani, P., Rudan, M.V., Liang, M., Medina-Rivera, A., Mohammed, H., Schmidt, D., Schwalie, P., Young, E.J., Reimand, J., Hadjur, S., Gingras, A.C. & Wilson, M.D. Topoisomerase II beta interacts with cohesin and CTCF at topological domain borders. Genome Biol 17, 182 (2016).

38. Bibel, M., Richter, J., Schrenk, K., Tucker, K.L., Staiger, V., Korte, M., Goetz, M. & Barde, Y.A. Differentiation of mouse embryonic stem cells into a defined neuronal lineage. Nat Neurosci 7, 1003–1009 (2004).

39. Tsumura, A., Hayakawa, T., Kumaki, Y., Takebayashi, S., Sakaue, M., Matsuoka, C., Shimotohno, K., Ishikawa, F., Li, E., Ueda, H.R., Nakayama, J. & Okano, M. Maintenance of self-renewal ability of mouse embryonic stem cells in the absence of DNA methyltransferases Dnmt1, Dnmt3a and Dnmt3b. Genes Cells 11, 805–814 (2006).

40. Thakurela, S., Garding, A., Jung, J., Schubeler, D., Burger, L. & Tiwari, V.K. Gene regulation and priming by topoisomerase IIalpha in embryonic stem cells. Nat Commun 4, 2478 (2013).

41. Ran, F.A., Hsu, P.D., Wright, J., Agarwala, V., Scott, D.A. & Zhang, F. Genome engineering using the CRISPR-Cas9 system. Nature protocols 8, 2281–2308 (2013).

42. Langmead, B. & Salzberg, S.L. Fast gapped-read alignment with Bowtie 2. Nat Methods 9, 357–359 (2012).

43. Ramirez, F., Dundar, F., Diehl, S., Gruning, B.A. & Manke, T. deepTools: a flexible platform for exploring deep-sequencing data. Nucleic Acids Res 42, W187–191 (2014).

44. Zang, C., Schones, D.E., Zeng, C., Cui, K., Zhao, K. & Peng, W. A clustering approach for identification of enriched domains from histone modification ChIP-Seq data. Bioinformatics 25, 1952–1958 (2009).

45. Jonathan A. Beagan, M.T.D., Katelyn R. Titus, Linda Zhou, Zhendong Cao, Jingjing Ma, Caroline V. Lachanski, Daniel R. Gillis, Jennifer E. Phillips-Cremins YY1 and CTCF orchestrate a 3-D chromatin looping switch during early neural lineage commitment. Genome Research (2017).

46. Zhang, Y., Liu, T., Meyer, C.A., Eeckhoute, J., Johnson, D.S., Bernstein, B.E., Nusbaum, C., Myers, R.M., Brown, M., Li, W. & Liu, X.S. Model-based analysis of ChIP-Seq (MACS). Genome Biol 9, R137 (2008).

47. Quinlan, A.R. & Hall, I.M. BEDTools: a flexible suite of utilities for comparing genomic features. Bioinformatics 26, 841–842 (2010).

48. Zhao, H., Sun, Z., Wang, J., Huang, H., Kocher, J.P. & Wang, L. CrossMap: a versatile tool for coordinate conversion between genome assemblies. Bioinformatics 30, 1006–1007 (2014).

49. Robinson, J.T., Thorvaldsdóttir, H., Winckler, W., Guttman, M., Lander, E.S., Getz, G. & Mesirov, J.P. Integrative genomics viewer. Nature Biotechnology (2011).

50. Liao, Y., Smyth, G.K. & Shi, W. The Subread aligner: fast, accurate and scalable read mapping by seed-and-vote. Nucleic Acids Res 41, e108 (2013).

51. Schmid, M.W. & Grossniklaus, U. Rcount: simple and flexible RNA-Seq read counting. Bioinformatics 31, 436–437 (2015).

52. Robinson, M.D. & Oshlack, A. A scaling normalization method for differential expression analysis of RNA-seq data. Genome Biology (2010).

53. de Dieuleveult, M., Yen, K., Hmitou, I., Depaux, A., Boussouar, F., Dargham, D.B., Jounier, S., Humbertclaude, H., Ribierre, F., Baulard, C., Farrell, N.P., Park, B., Keime, C., Carriere, L., Berlivet, S., Gut, M., Gut, I., Werner, M., Deleuze, J.F., Olaso, R., Aude, J.C., Chantalat, S., Pugh, B.F. & Gerard, M. Genome-wide nucleosome specificity and function of chromatin remodellers in ES cells. Nature (2016).

54. Huang, W., Sherman, B.T. & Lempicki, R.A. Systematic and integrative analysis of large gene lists using DAVID bioinformatics resources. Nat Protoc 4, 44–57 (2009).

55. Buenrostro, J.D., Giresi, P.G., Zaba, L.C., Chang, H.Y. & Greenleaf, W.J. Transposition of native chromatin for fast and sensitive epigenomic profiling of open chromatin, DNA-binding proteins and nucleosome position. Nat Methods 10, 1213–1218 (2013).

56. Lun, A.T. & Smyth, G.K. De novo detection of differentially bound regions for ChIP-seq data using peaks and windows: controlling error rates correctly. Nucleic Acids Res 42, e95 (2014).

57. Robinson, M.D., McCarthy, D.J. & Smyth, G.K. edgeR: a Bioconductor package for differential expression analysis of digital gene expression data. Bioinformatics 26, 139–140 (2010).

58. Yu, G., Wang, L.G. & He, Q.Y. ChIPseeker: an R/Bioconductor package for ChIP peak annotation, comparison and visualization. Bioinformatics 31, 2382–2383 (2015).

59. Li, H. & Durbin, R. Fast and accurate long-read alignment with Burrows-Wheeler transform. Bioinformatics 26, 589–595 (2010).

60. Wingett, S., Ewels, P., Furlan-Magaril, M., Nagano, T., Schoenfelder, S., Fraser, P. & Andrews, S. HiCUP: pipeline for mapping and processing Hi-C data. F1000Research 4, 1310 (2015).

61. Durand, N.C., Shamim, M.S., Machol, I., Rao, S.S., Huntley, M.H., Lander, E.S. & Aiden, E.L. Juicer Provides a One-Click System for Analyzing Loop-Resolution Hi-C Experiments. Cell systems 3, 95–98 (2016).

62. Hoffman, M.M., Ernst, J., Wilder, S.P., Kundaje, A., Harris, R.S., Libbrecht, M., Giardine, B., Ellenbogen, P.M., Bilmes, J.A., Birney, E., Hardison, R.C., Dunham, I., Kellis, M. & Noble, W.S. Integrative annotation of chromatin elements from ENCODE data. Nucleic acids research 41, 827–841 (2013).

63. Heger, A., Webber, C., Goodson, M., Ponting, C.P. & Lunter, G. GAT: a simulation framework for testing the association of genomic intervals. Bioinformatics 29, 2046–2048 (2013).

64. Beagan, J.A., Duong, M.T., Titus, K.R., Zhou, L., Cao, Z., Ma, J., Lachanski, C.V., Gillis, D.R. & Phillips-Cremins, J.E. YY1 and CTCF orchestrate a 3D chromatin looping switch during early neural lineage commitment. Genome Res 27, 1139–1152 (2017).

65. von Meyenn, F., Iurlaro, M., Habibi, E., Liu, N.Q., Salehzadeh-Yazdi, A., Santos, F., Petrini, E., Milagre, I., Yu, M., Xie, Z., Kroeze, L.I., Nesterova, T.B., Jansen, J.H., Xie, H., He, C., Reik, W. & Stunnenberg, H.G. Impairment of DNA Methylation Maintenance Is the Main Cause of Global Demethylation in Naive Embryonic Stem Cells. Mol Cell 62, 983 (2016).

66. Dykhuizen, E.C., Hargreaves, D.C., Miller, E.L., Cui, K., Korshunov, A., Kool, M., Pfister, S., Cho, Y.J., Zhao, K. & Crabtree, G.R. BAF complexes facilitate decatenation of DNA by topoisomerase IIalpha. Nature 497, 624–627 (2013).

67. Kagey, M.H., Newman, J.J., Bilodeau, S., Zhan, Y., Orlando, D.A., van Berkum, N.L., Ebmeier, C.C., Goossens, J., Rahl, P.B., Levine, S.S., Taatjes, D.J., Dekker, J. & Young, R.A. Mediator and cohesin connect gene expression and chromatin architecture. Nature 467, 430–435 (2010).

