## Supplementary Figures for "TIP5 safeguards genome architecture of ground-state pluripotent stem cells"

### Supplementary Figure 1

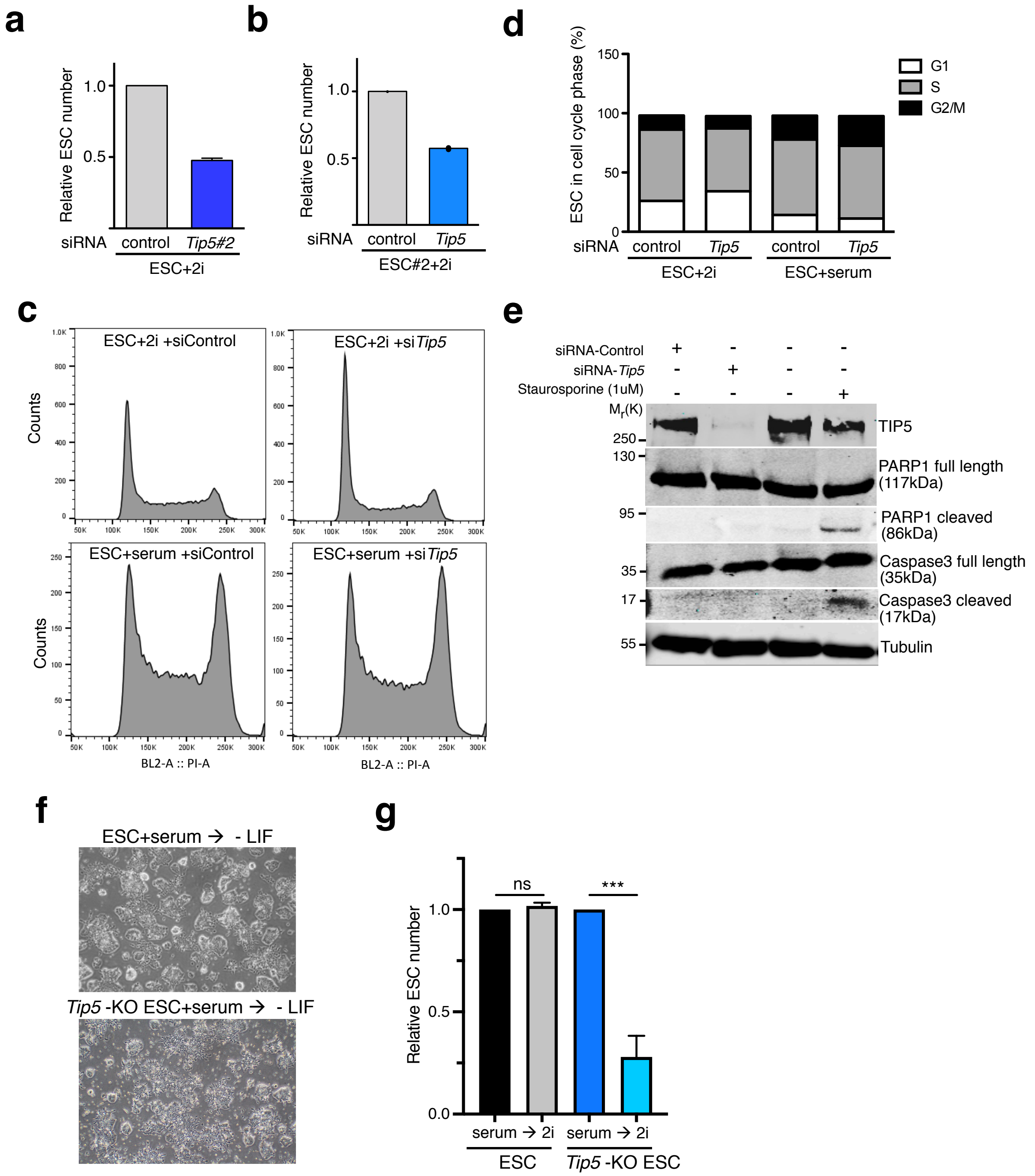

### Supplementary Figure 2

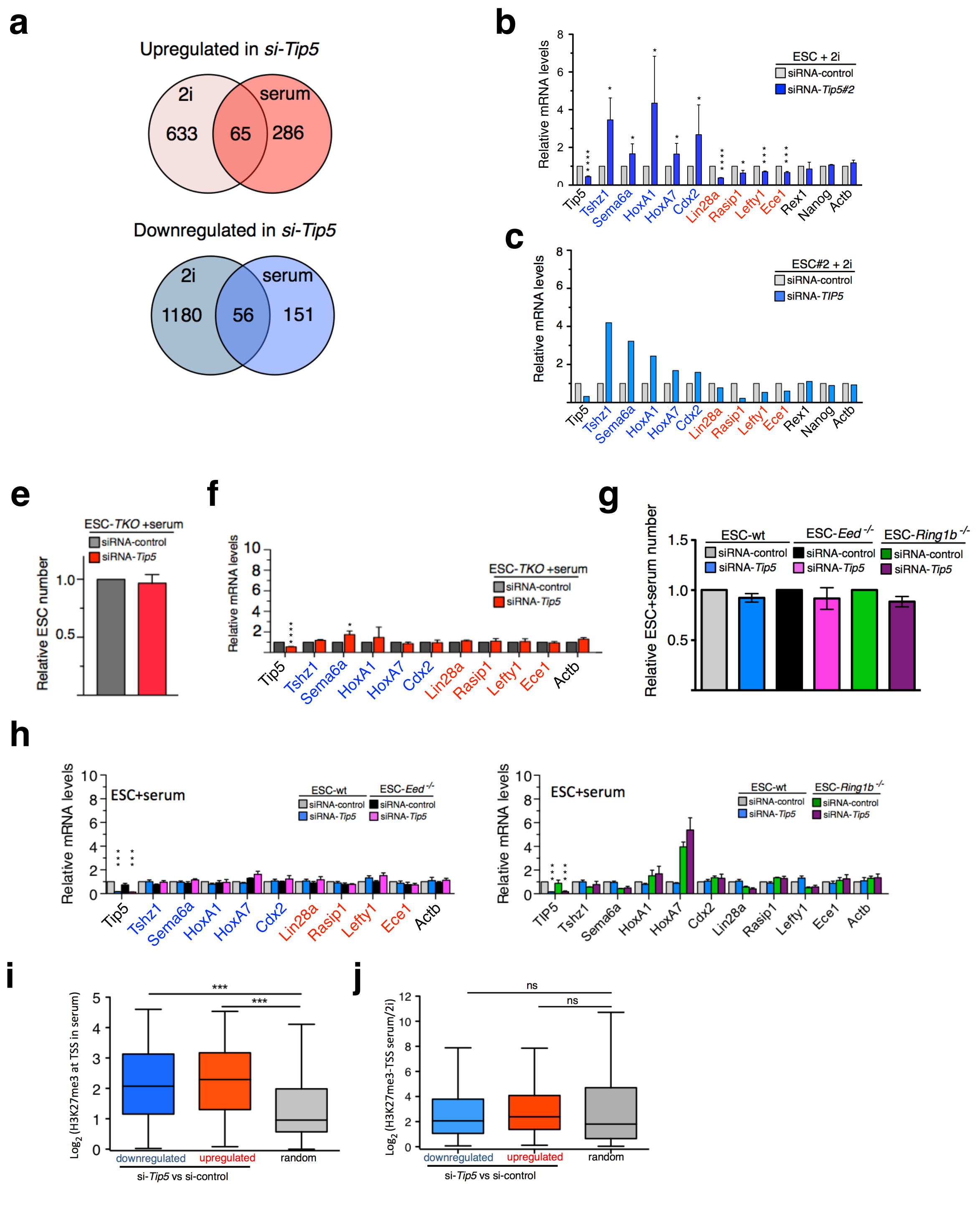

### Supplementary Figure 3

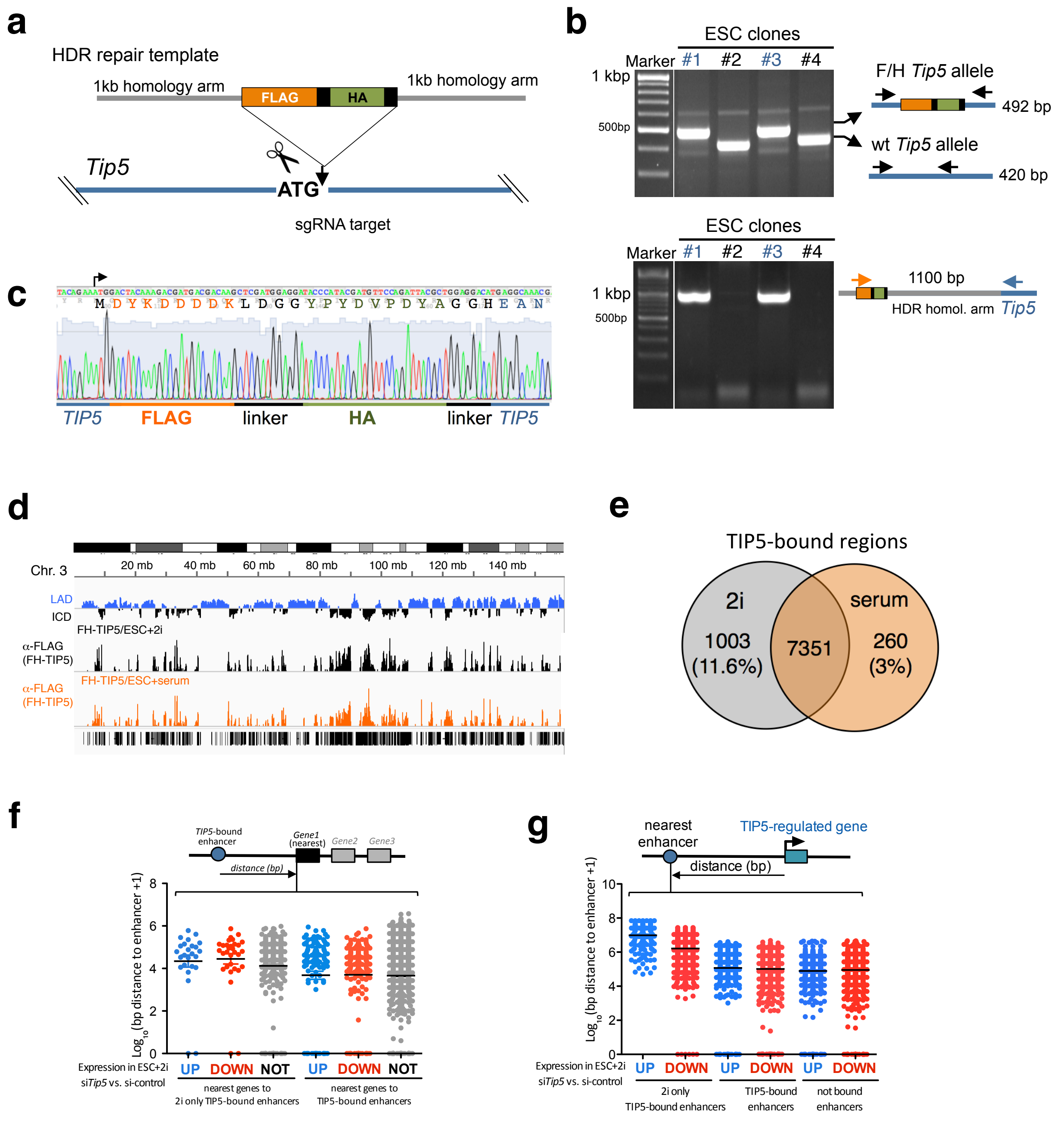

### Supplementary Figure 4

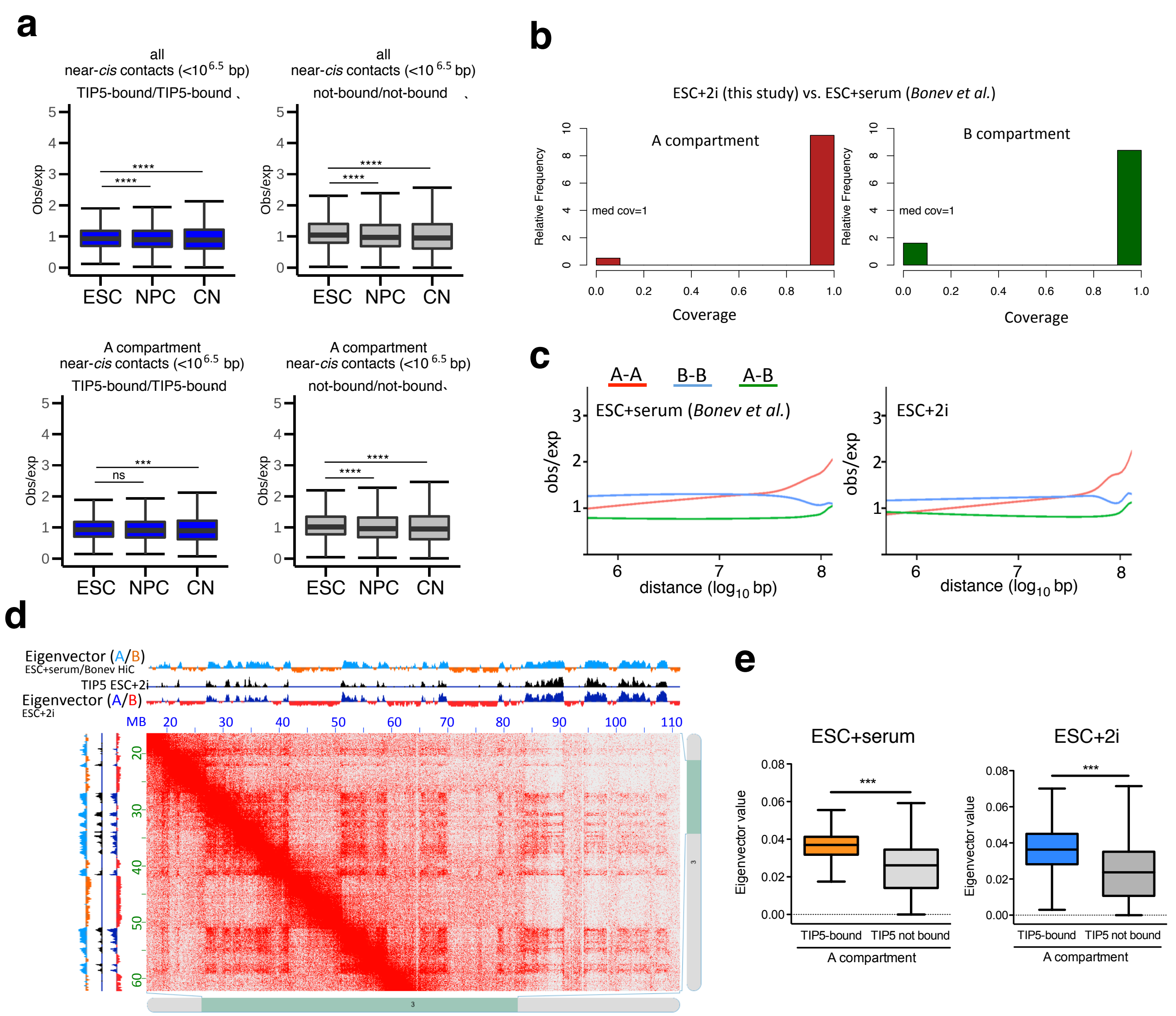

### Supplementary Figure 5

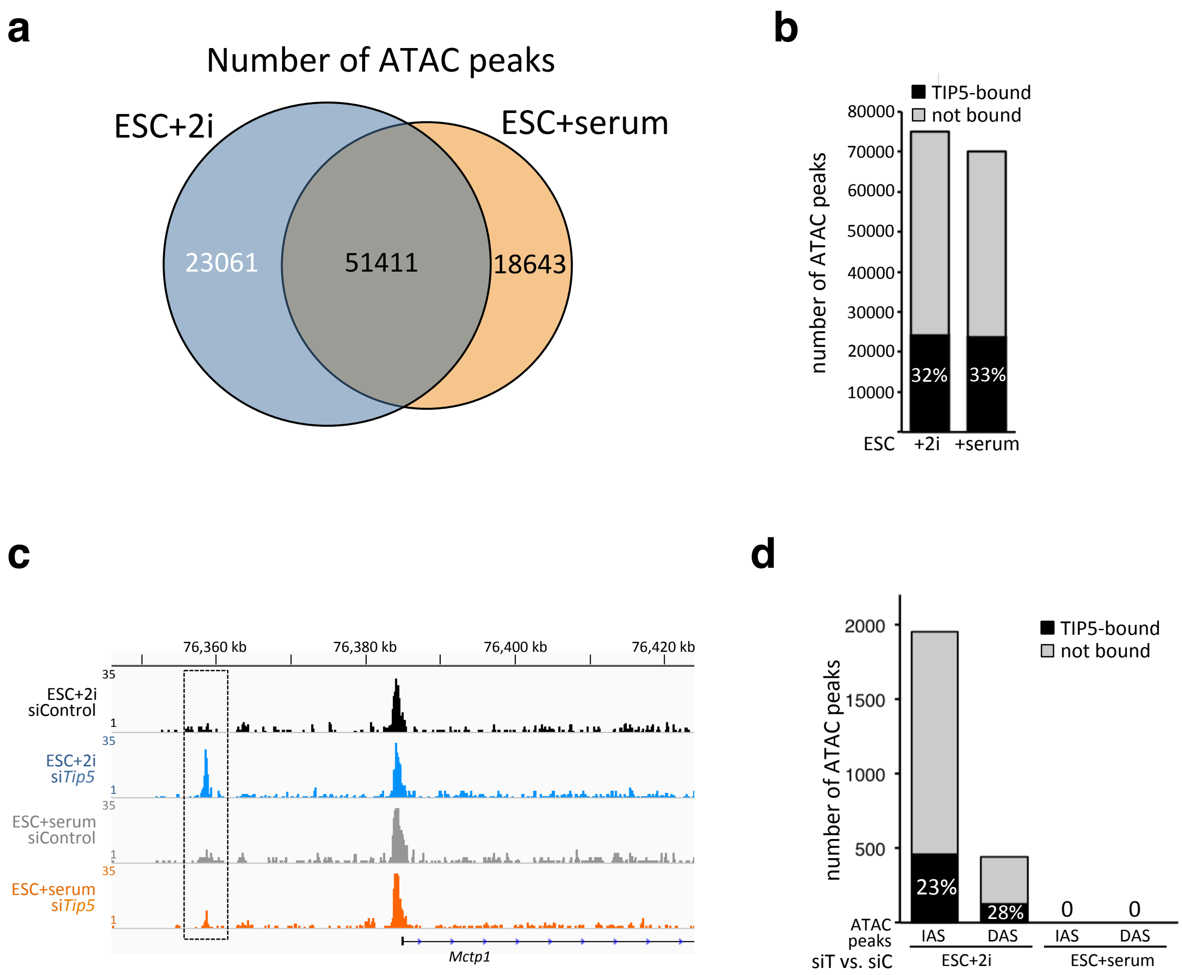

### Supplementary Figure 6

**a**

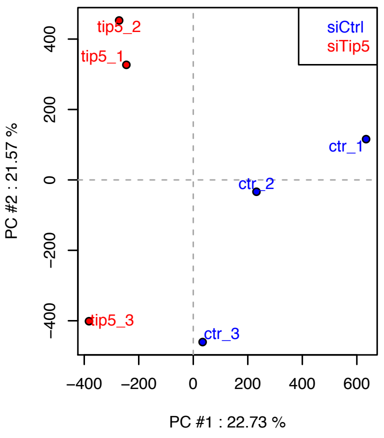


**b**

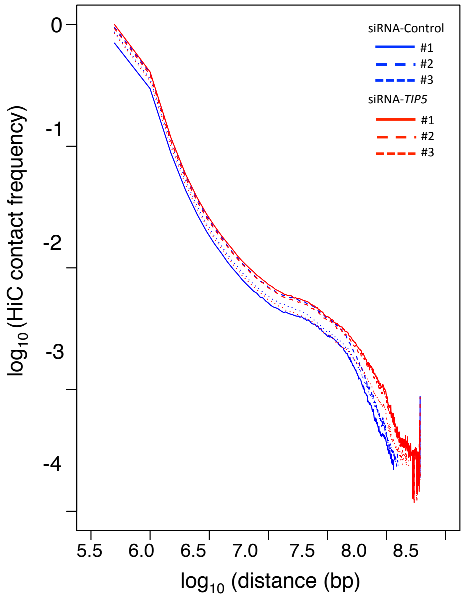


**c**

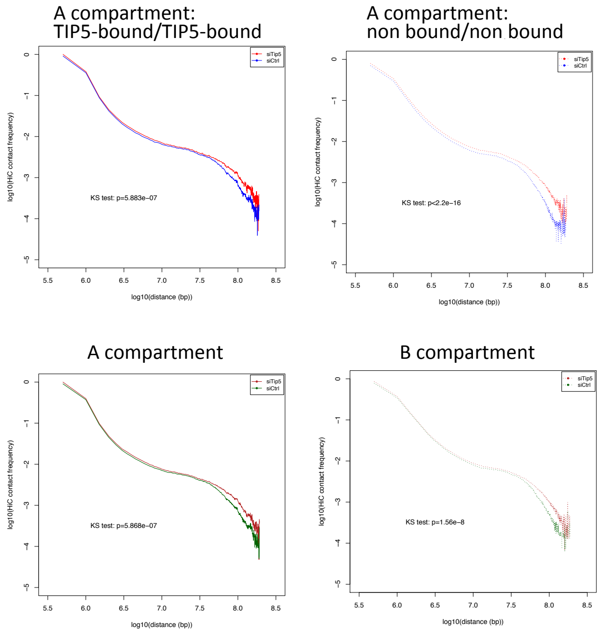

### Supplementary Figure 7

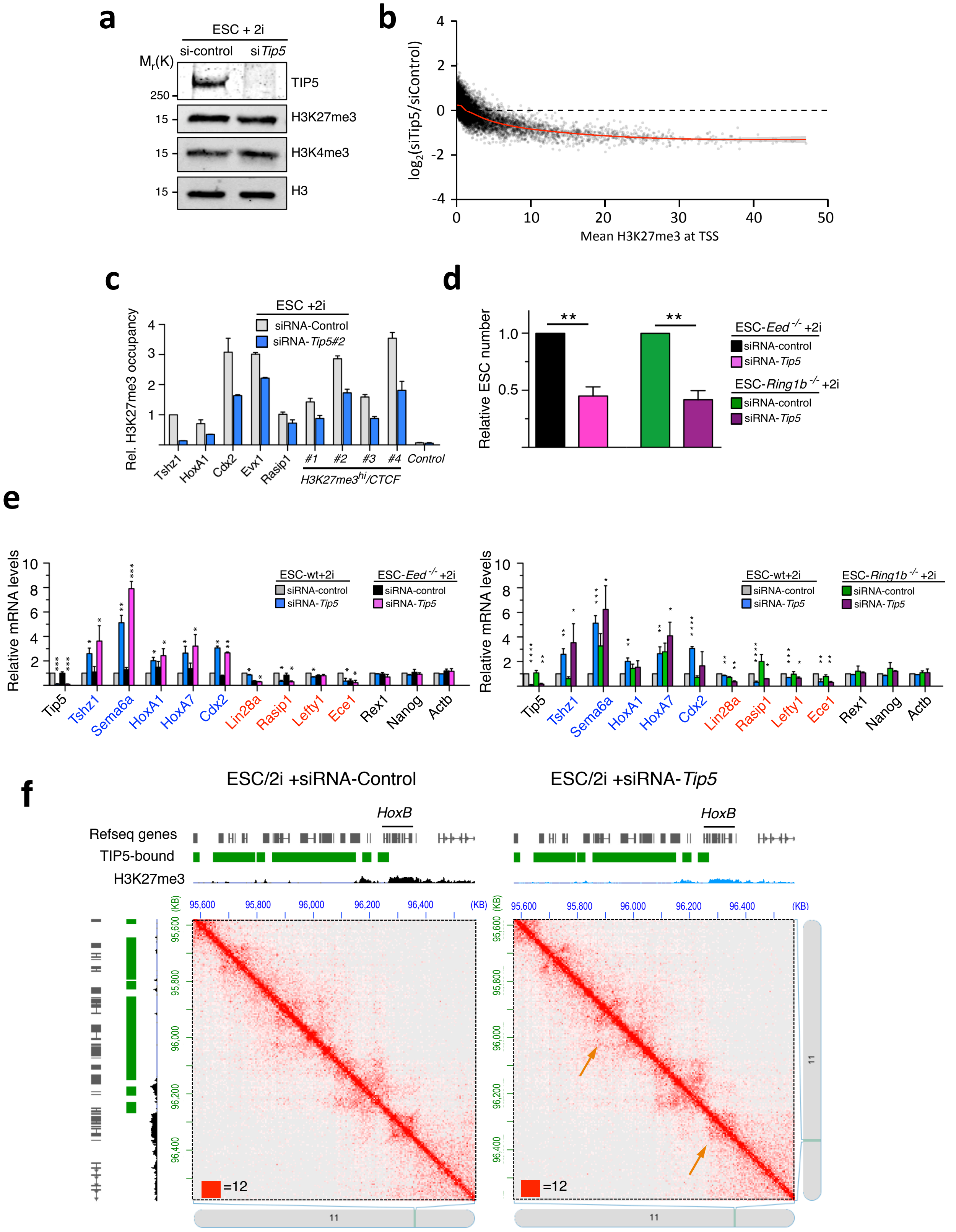

### Supplementary Figure 8

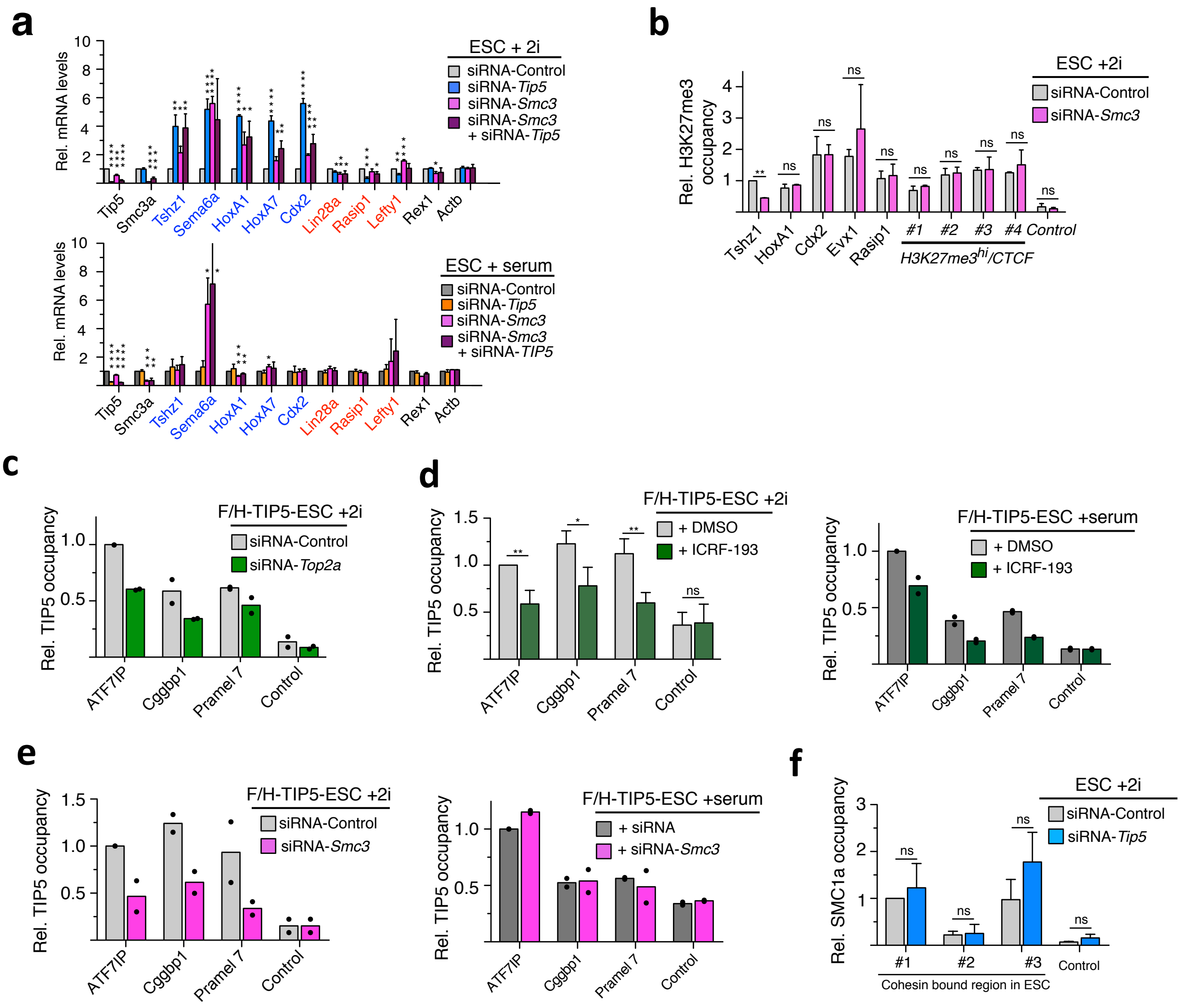
